# Genetic context effects can override canonical *cis* regulatory elements in *Escherichia coli*

**DOI:** 10.1101/2022.03.07.483377

**Authors:** Scott A. Scholz, Chase D. Lindeboom, Peter L. Freddolino

## Abstract

Recent experiments have shown that in addition to control by *cis* regulatory elements, the local chromosomal context of a gene also has a profound impact on its transcription. Although this chromosome-position dependent expression variation has been empirically mapped at high-resolution, the underlying causes of the variation have not been elucidated. Here, we demonstrate that 1 kb of flanking, non-coding synthetic sequences with a low frequency of guanosine and cytosine (GC) can dramatically reduce reporter expression compared to neutral and high GC-content flanks in *E. coli*. Despite the strong reduction in the maximal expression level from the fully-induced reporter, low GC synthetic flanks do not affect the time required to reach the maximal expression level after induction. Expression of the reporter construct is also affected by proximity to highly expressed ribosomal RNA operons depending on the relative orientation of transcription despite being insulated by strong transcriptional terminators, in a manner consistent with supercoiling competition. Overall, we demonstrate key determinants of transcriptional propensity that appear to act as tunable modulators of transcription, independent of regulatory sequences such as the promoter. These findings provide insight into the regulation of naturally occurring genes and specific rules for optimizing control of synthetic biology constructs.

## Introduction

Unicellular microbes,particularly fast-growing bacteria, typically have dense genomes with little intergenic DNA. Whereas variation in the transcribability of chromosomal regions in many eukaryotes is correlated with gene density, transcription from the *Escherichia coli* chromosome can vary many fold across regions of similarly high gene density (1). Several studies have reported variation in expression from a standardized reporter integrated into different locations in the *E. coli* genome (2–6). Several factors that affect the bacterial chromosome, some of which may also contribute to position-dependent expression variation, have been recently reviewed (7, 8). More recent studies have also examined the effects of metabolic burden and nutrient source on position-dependent expression variation (9). The variation in gene expression at different genomic loci in bacteria has been used to great effect in the field of synthetic biology to generate bacterial strains with improved production and circuit performance (10–15).

We recently generated an empirical, high-resolution map of transcription from a standardized reporter integrated at >144,000 sites in the *E. coli* genome. We termed the genomic position-dependent effects on reporter transcription “transcriptional propensity” (1). The chromosomal features that were observed to most strongly correlate with transcriptional propensity were local GC content, proximity to ribosomal RNA operons, and binding of the nucleoid associated proteins (NAPs) H-NS and Fis. For example, H-NS binding sites are apparent as troughs on the transcriptional propensity map (i.e., low transcriptional propensity). Subsequent high-throughput assessment of fluorescent protein expression from transposon integrations largely confirmed these results, and also linked H-NS binding sites to low reporter expression (16). H-NS binds to and reduces expression of genes with AT rich sequences (17–20). H-NS alone and in combination with other factors has been shown to function to drive RNAP pausing after transcription initiation (21). In our previous work, we observed that reporters that integrated into H-NS binding regions also became silenced, despite the fact that the GC-neutral reporter sequence was unaltered.

At the other extreme, the most readily apparent features of transcriptional propensity topology are broad peaks of high transcriptional propensity that are centered on ribosomal RNA operons (*rrn*; see **Supplementary Fig. 1A**). *rrn* are the most heavily transcribed genes in the *E. coli* genome (22, 23), and additional highly-transcribed genes (including elements of the amino acid biosynthetic pathways) are enriched in regions of high transcription propensity around *rrn* (Scholz et al. 2019). A model in which *rrn* colocalize into nucleolus-like domains (24, 25) and thereby increase RNAP access to the surrounding genes is consistent with this data and may explain the transcriptional propensity peaks at *rrn*, but requires further experimental testing.

Context-dependent expression variation is also caused by interference between transcription of neighboring genes, as has been demonstrated through rational reporter integration and construct design (3, 26–29). In general, terminator-flanked genes downstream of highly expressed promoters are depressed in their expression, consistent with a build-up of positive supercoiling. Supercoiling interference effects are also apparent for genes encoded in the divergent orientation (26, 30).

In this study, we investigated the relative contributions of local GC content, the presence of proximal ribosomal RNA operons, and supercoiling competition on expression of a transcriptionally insulated reporter. Consistent with prior work, reporter expression varied substantially depending on the chromosomal integration site and our genetic context manipulations. We found that a reporter is silenced if it is flanked by 1 kb of non-coding low GC content DNA, while high GC flanks can cause de-repression in a context that would otherwise be silencing. Despite the substantial differences in reporter expression depending on the integration site and the GC content of flanking DNA, the time by which the reporter reaches maximal fluorescence level after induction is similar. We also demonstrate that although *rrn* are at the center of large transcriptional propensity peaks, removal of the *rrn* itself does not diminish the expression of a reporter that has integrated in close proximity to it. Additionally, the transcriptionally insulated reporter used in this study was affected by interference from the highly expressed *rrnE* operon despite an absence of direct transcriptional readthrough. This finding is consistent with previously observed supercoiling competition effects. Genetic context effects, reported throughout this article at the population and single-cell level, appear to cause tunable and uniform changes in expression across a clonal population, as opposed to multimodal effects on expression.

## Materials and Methods

### Strains and media

All strains were constructed from ecSAS17, a derivative of *E. coli* K12 MG1655 with chromosomal integrations of constitutively expressed tet repressor and mCherry (1). The mNeonGreen reporter construct was chromosomally integrated into different genomic locations with or without synthetic genetic contexts, but otherwise unaltered from our previous study. For strain construction, ecSAS17 harboring the lambda Red plasmid pSIM5 (gift from Prof. Don Court) was transformed with linear PCR fragments of the reporter or reporter flanked by synthetic GC content (sequences of the plasmids used for strain construction are given in **Supplementary Data 1**, and a full listing of strains used in the present study is given in **Supplementary Data 2**). The reporter kanamycin resistance marker was then removed from each strain with the helper plasmid pCP20. Targeted integration of the reporter construct, with or without flanking synthetic genetic contexts as indicated, was confirmed by PCR genotyping across integration junctions and sanger sequencing of junction PCR products (a complete listing of primers is given in **Supplementary Data 3**). For all strain construction, LB lennox broth with the appropriate antibiotics added was used for propagation and growth. Confirmed strains were cryopreserved indefinitely at -80°C in 15% glycerol.

### Strain growth for microplate reader and flow cytometry strain assessment

Each strain was independently streaked onto an LB agar plate from a cryopreserved stock and incubated overnight at 37°C. Plates were stored for up to 12 days at 4°C. A single colony from the LB agar plate was inoculated into 3 mL of LB Lennox and grown at 37°C overnight with shaking. 10 μl of the overnight culture was inoculated into 1 mL M9RDM (Glucose 4 g/L, NH4Cl 1 g/L, KH2PO4 3 g/L, NaCl 0.5 g/L, Na2HPO4 6 g/L, MgSO4 240.7 mg/L, ferric citrate 2.45 mg/L, CaCl2 111 ng/L, 200 mL/L 5x Supplement EZ, 100 mL/L 10X ACGU (Teknova cat # M2103), 1 mL/L micronutrient solution) + a final concentration of 100 mg/L anhydrotetracycline (aTc) and incubated at 37°C for 2 hours with shaking. 1 μl of the log-phase cells was inoculated into 150 μl of M9RDM + aTc in a microplate (96-well flat clear bottom, black sides or flat bottom all clear for microplate reader and flow cytometry, respectively) and topped with 100 μl mineral oil. For microplate reader assessment, mNeonGreen fluorescence (Ex. 506 nm, Em. 550 nm), mCherry fluorescence (Ex. 580 nm, Em. 610 nm) and optical density (OD) at 450 nm were measured every 10 minutes. For growth before flow cytometry assessment OD 450 was measured every 10 minutes. All strains were otherwise grown at 37°C with orbital shaking in a Synergy H1 or Epoch BioTek plate reader. For flow cytometry assessment, the microplate was rapidly moved from the microplate reader into an ice water bath after 3 hours of growth and immediately transferred for auto-sampled flow cytometry.

### Flow cytometry and single-cell analysis

Rapidly cooled cells grown to log phase were sampled from an ice-cold 96-well plate on a Miltenyi Biotec MACSQuant YVB flow cytometer cooling auto-sampler. After daily calibration with MACSQuant Calibration Beads, forward scatter (FSC), side scatter (SSC), mCherry (Ex. 561 nm, Em. band pass 615/20 nm) and mNeonGreen (Ex. 488 nm, Em. band pass 525/50) signals were collected with an automated sampling and washing procedure. We calculated the correlation between the mNG signal and the signals for FSC, SSC and mCherry, as well as the slope of the linear regression of log-transformed values with respect to log-transformed mNG within each strain. Since the correlation between mNG and SSC was the strongest, we corrected log mNG for each strain by subtracting the product of the average slope of log mNG with respect to log SSC for each cell in order to calculate mNG concentration. This correction tightens the distribution of mNG signal within each strain by normalizing the natural variation attributable to cell morphology.

### Microplate reader data analysis for cells growing in log phase

All population-level fluorescence per OD values presented in bar-plots in this article are the geometric mean of fluorescence per OD over all time points in which OD450 was between 0.15 to 0.35, which is within the log growth phase for all strains. The fluorescence per OD values from each replicate are also shown as dots on every bar plot. The autofluorescence per OD of the reporter-free base strain ecSAS17 calculated over the same OD range (grown in parallel within the same microplate) was subtracted from the fluorescence per OD value from each strain.

### Induction timing of strains passaged in a microplate reader

Since the autofluorescence per OD from reporter-free cells growing in M9RDM medium changes at different ODs, we performed a kernel regression of the autofluorescence signal of ecSAS17 cells growing in M9RDM with respect to the optical density at 450 nm (OD 450) in order to calculate the fluorescence signal attributable to the growth medium and cells at every OD. As the reporter-free cells grow in M9RDM, the fluorescence signal initially dips slightly but substantially (presumably due to consumption of a medium component contributing to autofluorescence signal, as this decrease was not observed in cell-free medium) before increasing as a result of autofluorescence of the cells. Therefore, we subtracted the expected autofluorescence signal from cells growing in M9RDM derived from the kernel regression from the fluorescence signal from each strain at every measured OD value and finally divided by OD in order to report only the fluorescence per OD resulting from mNG reporter expression itself in **Figure 3B**. The fold-difference between fluorescence per OD and maximal induction level was calculated by averaging the fluorescence per OD between minute 410 and 530 minutes and reporting the log2 fold-change of the fluorescence per OD at each timepoint from that maximal induction value of each strain.

### qPCR based reporter measurements

The calibration of reporter expression by RT-qPCR shown in **Supplementary Figure 2** was performed as described in (1). In brief, cells were grown to OD 450 0.2 in M9RDM, at which point 650 μl of the culture was mixed with 1.3 mL RNAProtect Bacterial Reagent (Qiagen) and frozen according to the manufacturer’s instructions. RNA was then purified using the RNeasy mini kit (Qiagen). An additional DNaseI treatment (Qiagen RNase-Free DNase Set) was performed on the purified sample in order to remove residual gDNA and re-purified using the RNeasy kit. cDNA was finally synthesized using the NEB Protoscript II First Strand cDNA synthesis kit (New England Biolabs) following the manufacturer’s instructions in parallel to control samples without the Reverse Transcriptase enzyme (-RT).

qPCR reactions on the cDNA and -RT control samples described above were performed with primers for mNG 63-64 and *mdoG* primers 65-66 (see **Supplementary Data 3**) using BioRad iTaq Universal SYBR Green Supermix, on a BioRad CFX96 instrument. In all cases the -RT samples were at least 6 Ct higher than the matched cDNA sample (comparable to water-only controls). In each case, we calculated the expression levels based on the Ct values obtained for the mNG reporter in the cDNA sample normalized by the signal from mdoG.

## Results

### Genome context has similar effects on reporters with different promoter strengths

To study the impact of genetic context on gene expression, we measured expression from a fluorescent reporter construct integrated at several genomic loci in the *E. coli* genome with manipulated local genetic contexts. As detailed in **Supplementary Text 1**, we found that translated protein levels in our reporter serve as a faithful readout of transcriptional propensity across a wide range of expression levels, both in bulk (plate reader) and at a single cell level (flow cytometry). Thus, for the sake of convenience, we made use of fluorescent readouts of protein production as a proxy for the context-dependent transcriptional changes in reporter expression in the work described here. An important corollary of the data shown in **Supplementary Text 1** is that we see no evidence for any position-dependent “translational propensity” in *E. coli* (at least for the limited set of sites considered here). If the translation of an mRNA depends upon its location on the chromosome, we would expect to see position-dependent differences in the protein:RNA ratio; instead, fluorescent protein levels are well correlated both with transcriptional propensity and RNA levels (**Supplementary Fig. 2**).

As the targeted reporter integrations using the TetO1 promoter from our prior work are generally expressed as expected based on the transcriptional propensity map (1) (**Supplementary Fig. 1B-C**; see also **Supplementary Text 1**), we sought to understand whether different promoters driving reporter expression would be similarly affected by genetic context. To test this, we integrated reporters with four promoter variants into three genetic loci that are proximal to each other (maximum distance is 76 kbp or <2% of total genome length apart, to minimize dosage effects), but have very different transcriptional propensity (*yagF, eaeH*, and *yafT*; see **Supplementary Fig. 1**). Throughout this work, the name assigned to an insertion location refers to the closest gene with a coordinate number preceding reporter integrations. We observed via bulk fluorescence experiments that mNG expression driven by the same TetO1 promoter used to generate the transcriptional propensity map ranked as expected. mNG expression driven by different promoters was affected by genetic context in the same rank order (*yagF* > *eaeH* > *yafT*), unless very weak promoters are used, in which case the distributions become essentially indistinguishable (**Fig. 1A-B)**. The strains with the weakest promoter that we tested still produced a fluorescent signal above the reporter-free base strain autofluorescence level, but did not change based on genetic context, indicating there may be a baseline level of transcriptional activity that cannot be further silenced by local genetic context. We observed a similar rank ordering in flow cytometry experiments for the TetO1 and *rplM* promoters, although in this case the distinctions between locations for the weaker *gmk* and *yghU* promoters were not apparent (**Fig. 1C**). Flow cytometry may lack the sensitivity to reliably detect such differences at the extreme low end of expression level, especially given that a large fraction of the cells with the *gmk* and *yghU* promoters had fluorescence levels below the average for reporter-free cells (**Fig. 1C**). Importantly, the flow cytometry results demonstrate that the contexts we have tested here only shift the fluorescence of cells as a unimodal population, with no instances of the appearance of a secondary population; thus, the effects of chromosomal context represent tunable, rather than all-or-nothing, changes in expression.

**Figure 1:**
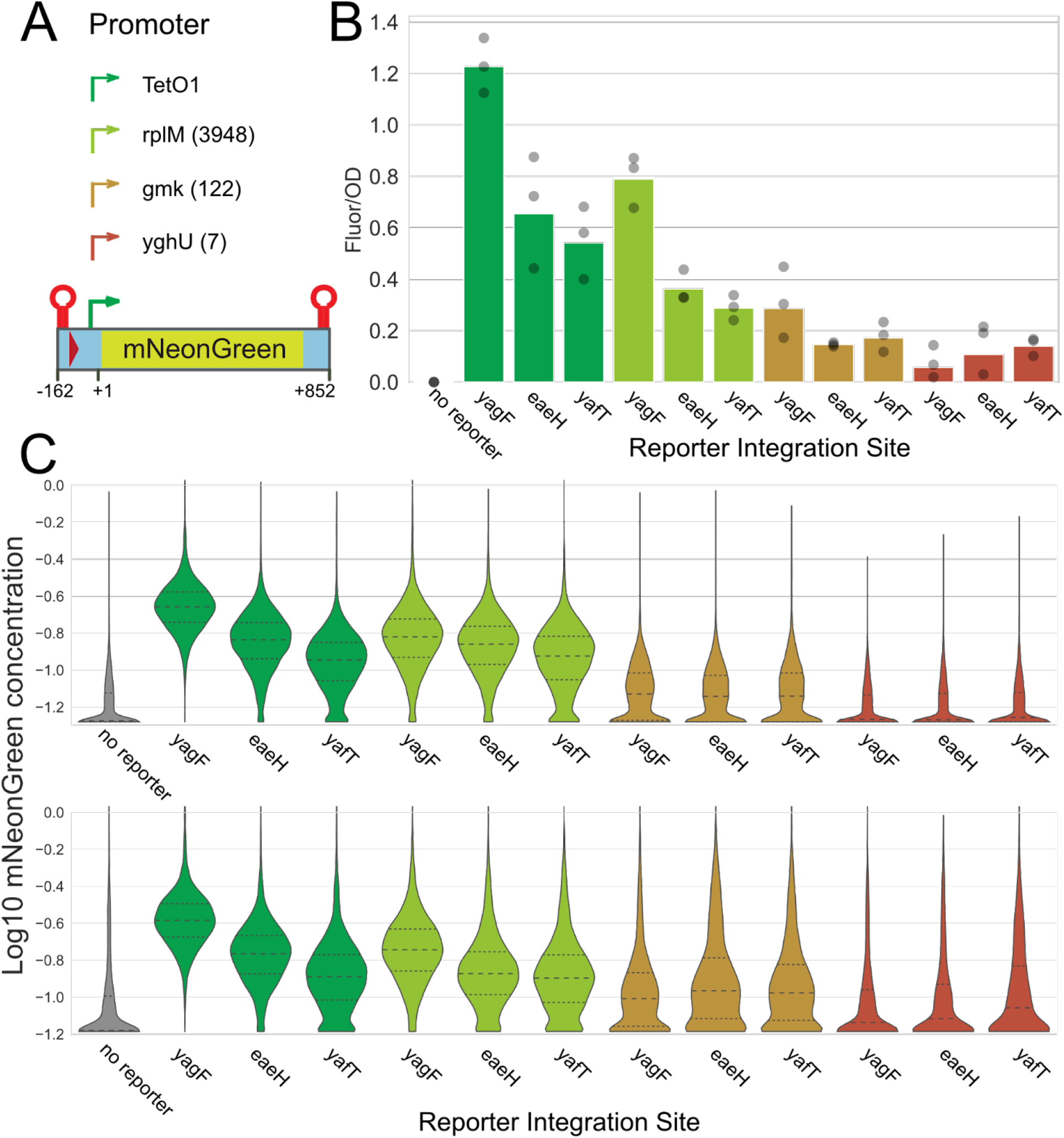
Genetic context affects gene expression from strong and weak promoters. A) Diagram of the mNeonGreen reporter with different strength promoters, including the transcripts per million normalized RNA count from the native gene in parentheses (31). Coordinates for the reporter construct diagram are relative to the transcription start site of the TetO1 promoter. B) Microplate reader normalized Fluorescence/OD for mNeonGreen reporters with four different promoters integrated into three adjacent genomic loci (see **Supplementary Fig. 1A**). Bars show the geometric mean signal across three biological replicates, with individual replicates shown as grey circles. C) Flow cytometry readings of the same strains measured on two different days show that the distribution of mNeonGreen concentration per cell is consistent with microplate data. All cells with fluorescence below the median for the ‘no reporter’ cells are clamped at a lower limit equal to the ‘no reporter’ median, representing the limit of detection in the experiment; the apparent accumulation of cells at the bottom of the axis this does not represent a secondary definable population, but rather, simply the presence of a portion of cells that do not have detectable fluorescence. The dashed line shows the median, and dotted lines 25th/75th percentile.

### Synthetic genetic context can strongly influence reporter expression

Transcriptional propensity is strongly correlated with genomic features such as local GC content, binding of the nucleoid associated protein H-NS, and proximity to ribosomal RNA operons (1). We set out to directly test whether systemic alteration of the most informative of these genetic features could affect expression of an integrated reporter. In all cases, our standard reporter with a TetO1 promoter was used (and as shown in **Figure 1A**, the sequence context was fixed and constant across all integrations for 162 bp upstream of the transcription start site). An identical reporter construct was integrated into two sites, one with naturally high (*nfi*) and one with naturally low (*wbbH*) transcriptional propensity. In order to isolate the effect of flanking GC content on reporter expression, we then integrated the same reporter into both sites and included 1 kb of flanking synthetic context on each side with either 35% average GC content or 65% average GC content. These GC contents represent extremes compared to the average *E. coli* GC content of 50.4% (**Fig. 2A**). We amplified two independent sequences for both our 35% and 65% GC synthetic flanks from eukaryotic yeast or human DNA (“35% GC” A/B and “65% GC” A/B respectively), which are not expected to bear specific regulatory elements that are functional in *E. coli*. All sequences are given in **Supplementary Data 1**.

**Figure 2:**
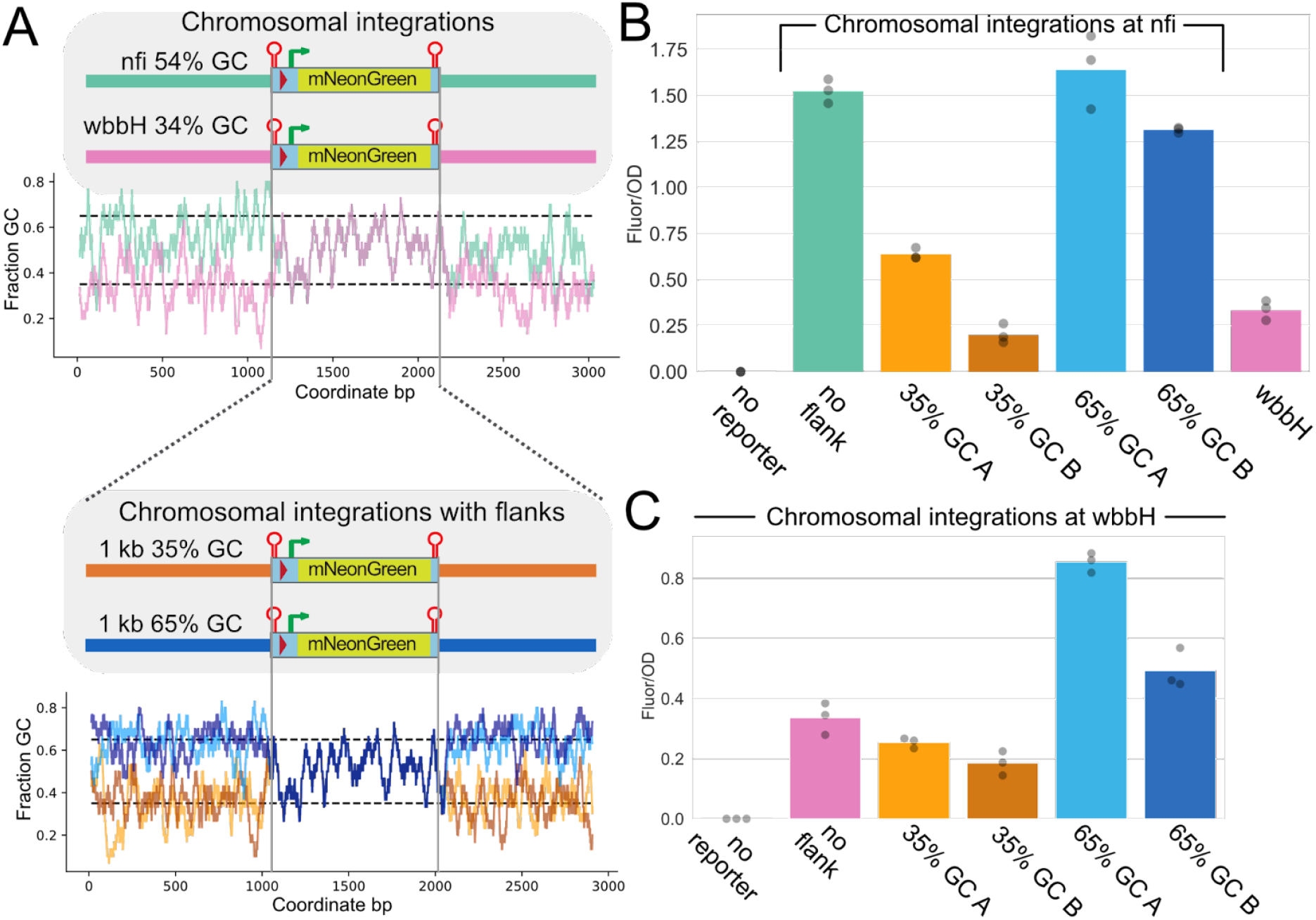
Local low GC content silences reporter gene expression at two sites. A) Diagram of the mNG reporter integrated into the *nfi* (Teal) or *wbbH* (Pink) locus or the construct flanked by synthetic genetic contexts of 35% (Orange) or 65% (Blue) GC content and integrated into either the *nfi* or *wbbH* genomic locus. For the synthetic flanks, the light and dark colors represent the two different variants that were used (and correspond to colors in the following panels). The accompanying plot shows GC fraction over a 30 bp sliding window of the mNG construct integrated into the genome (top) or into the genome with synthetic flanks (bottom). 35% GC content is the 1.67 percentile and 65% GC content is the 99.88 percentile of all 500 bp sliding windows of GC content. B) Reporter fluorescence per OD 450 measured in three independent microplate reader experiments for reporters integrated at the *nfi* locus. A chromosomal integration without synthetic flanks at *nfi* and *wbbH* is included for reference. Bars show the geometric means of three biological replicates (which are themselves shown as grey dots). C) Reporter fluorescence per OD 450 measured in three independent microplate reader experiments for reporters integrated at the *wbbH* locus. Bars show the geometric means of three biological replicates (which are themselves shown as gray dots).

Fluorescence from the reporter expressed from within low GC content synthetic contexts was a fraction of the reporter with no synthetic flanks, regardless of the intrinsic transcriptional propensity of the integration site itself, as shown in **Figure 2B-C**.The addition of low-GC flanking regions substantially decreased expression of the reporter at both the *nfi* and *wbbH* loci, with the effects more dramatically evident at *nfi* due to the initially high expression there (**Fig. 2B-C**). Compared to the flank-free reporter, high GC content synthetic flanks did not have a strong impact on reporter expression when the reporter construct was integrated at the high transcriptional propensity site (**Fig. 2B**). However, they substantially increased fluorescence at the low transcriptional propensity site (**Fig. 2C**). We observed consistent results via flow cytometry, and again observed tunable changes in expression in response to the changing GC content of the flanking regions (**Supplementary Figs. 3** and **4;** note that mCherry is always at the same site as part of the base strain and is integrated near *yihG*). Likewise, we also observed that synthetic flanks have strong effects on mNG expression from the same reporter encoded on a plasmid, with 65% GC flanks substantially increasing fluorescent reporter expression compared with 35% GC flanks (see **Supplementary Fig. 5**). Together, our findings suggest that the addition of ∼1kb flanks with extreme GC content can substantially alter the transcriptional propensity of a site even when the promoter region itself is held fixed. However, GC content cannot be the sole factor leading to position-dependent transcriptional propensity given that altered flanking sequence composition cannot reliably convert the propensity at *wbbH* to that at *nfi* (at the very least, dosage effects (32, 33) almost undoubtedly make orthogonal contributions, and other factors may as well).

### Context-driven variations in reporter expression affect equilibrium expression levels but not induction kinetics

Due to the experimental setup of our high-throughput transcriptional propensity profiling, which involved induction for a fixed period of time prior to measurement of transcript levels, low measured transcriptional propensity could arise due to either slow induction of transcription, or low equilibrium levels of transcription at those sites. In principle, It is possible that measured low propensity sites may take longer to reach maximal expression level and might eventually match the fluorescence level of cells with reporters at high-propensity sites after additional cell doublings in inducing conditions.

To assess the induction rates and maximal expression during induction in the context of our engineered high- and low-propensity reporters (the high- and low-GC flanked reporters discussed in the context of **Figure 2**, plus a medium propensity unflanked reporter at *eaeH*, in all cases under control of the TetO1 promoter), we passaged growing cells from M9RDM into M9RDM + anhydrotetracycline (aTc) with additional subsequent passages into the induction medium in a microplate to maintain the cells in exponential growth. Fluorescence and optical density at 450 nm were monitored every 10 minutes (**Fig. 3A**; induction begins at 210 minutes in the timeline on that figure). The maximal fluorescence reached by cells with the reporter integrated at three different sites with different synthetic flanks varied greatly even after multiple passages in the induction medium. Fluorescence levels were consistent with that observed in microplate reader and flow cytometry measurements from previous experiments (e.g., **Figure 2**), and most notably, the equilibrium fluorescence level achieved after several passages – and retained after stationary phase entry – was strongly determined by the transcriptional propensity of the reporter site (**Fig. 3B**). The time required to reach maximal fluorescence was very similar for all strains (**Fig. 3C**), typically stabilizing within 120 minutes of induction (noting the lag before the observed differences between the observed and maximum values stabilized near 0). Tracking fluorescence at the lower end of cell density after culture passaging introduced enough technical noise to make precise calculations of time required to reach maximal induction unreliable, as transient spikes/dips in the OD-normalized mNeonGreen signal are observed at low OD after dilution. Nevertheless, we can assert that variations in induction timing, if any, are on the order of tens of minutes (**Fig. 3C**). For comparison, the mixed reporter library strain that was used to generate our previous high-resolution transcriptional propensity map was induced for 210 minutes prior to harvest (1), equivalent to the 420th minute in the induction experiment, which corresponds well to the equilibrium expression level. Thus, we find that the expression levels reported throughout this work and in (1) are comparably in the log-phase of growth and correspond to fully induced time-points (**Fig. 3**).

**Figure 3:**
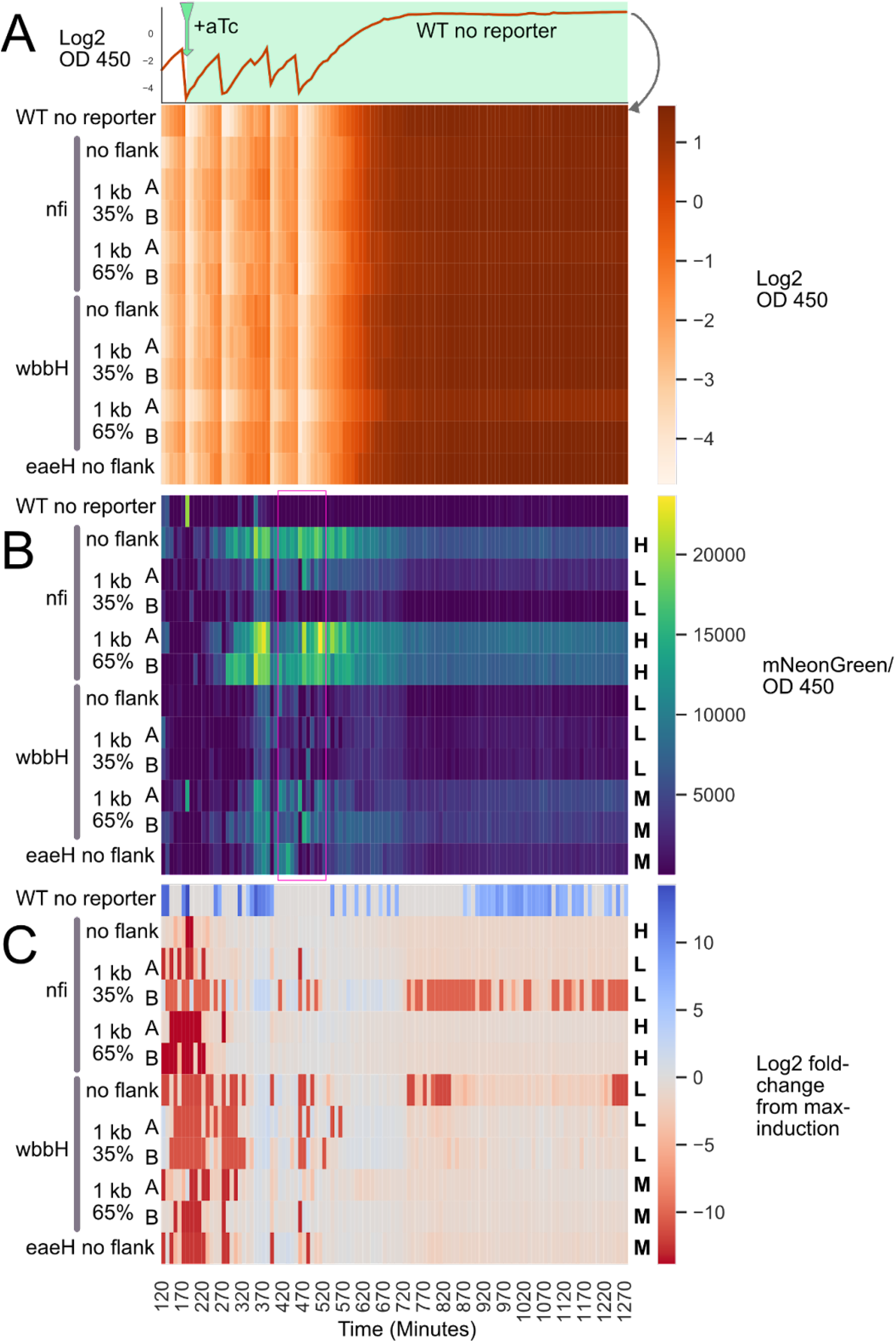
Genetic context effects are robust across timepoints. 12 strains, with and without synthetic flanks, were grown and passaged into fresh medium during growth in a microplate reader. The inducer aTc was added at the first passage time (Green arrow, 210 minutes) and included in every subsequent passage. A reporter integration at the *eaeH* site was included as a “medium” expression site for comparison (1 kb of flanking sequence has a GC content of 37%). A) The heat map coloration indicates the log2 of OD measured at 450 nm. The data for the log2 OD 450 for only the WT strain without a reporter is shown in a line plot directly above the heat map as a clarifying example representation of the heatmap data. B) Fluorescence per OD 450 for all 12 strains. The H, M, and L letters stand for high, medium and low expression, respectively, expectations as measured from traditional microplate reader experiments (without passaging). C) Log2 fold-change from maximal induction (Defined by the average Fluor/OD included in the Purple Box in panel B) shows that induction timing from reporter strains with different synthetic genetic contexts is similar.

### Ribosomal RNA operons themselves are not necessary for ribosomal RNA operon-proximal transcriptional propensity peaks

Because several transcriptional propensity peaks are centered on ribosomal RNA operons (*rrn*; see **Fig. S1A**), we sought to determine whether the *rrn* operons cause the peak formation. To that end, we either integrated our standard reporter construct upstream of rrn promoters and their associated Fis binding sites (*rrn* int, in which the Fis sites are unperturbed), or replaced the entire operon and its promoter region with the reporter (*rrn* KO; see **Fig. 4**). Despite the striking centrality of rrn in transcriptional propensity peaks, replacing an *rrn* with the reporter (*rrn* KO) did not cause any decrease in reporter expression compared to the upstream integration. Additionally, we did not detect any changes in growth rate (measured via plate reader growth curves) or overall cell protein expression (using our internal mCherry control as measured by flow cytometry), either of which could otherwise confound interpretation of fluorescence data if they changed (**Supplementary Figs. 6** and **7**). Thus, we find that the effects of proximity to the *rrn* are completely independent of the presence of *rrn* transcription or even the *rrn* operon itself, and must instead reflect some more general characteristic of the chromosomal regions in which *rrn* operons are found. To further investigate the roles played by different portions of the *rrn*-proximal regions in setting the transcriptional propensity near them, we also tested a third *rrnE* variant in which the ribosomal operon and its annotated promoters were removed, but the upstream Fis binding sites (known to be important in regulating transcription) were retained -- we refer to this construct as “KO+”. The “KO+” variant showed a modest increase in fluorescence even over the corresponding “KO”, suggesting that the Fis binding sites may yet further increase RNA polymerase recruitment to this region even in the absence of the *rrn* operon itself.

**Figure 4:**
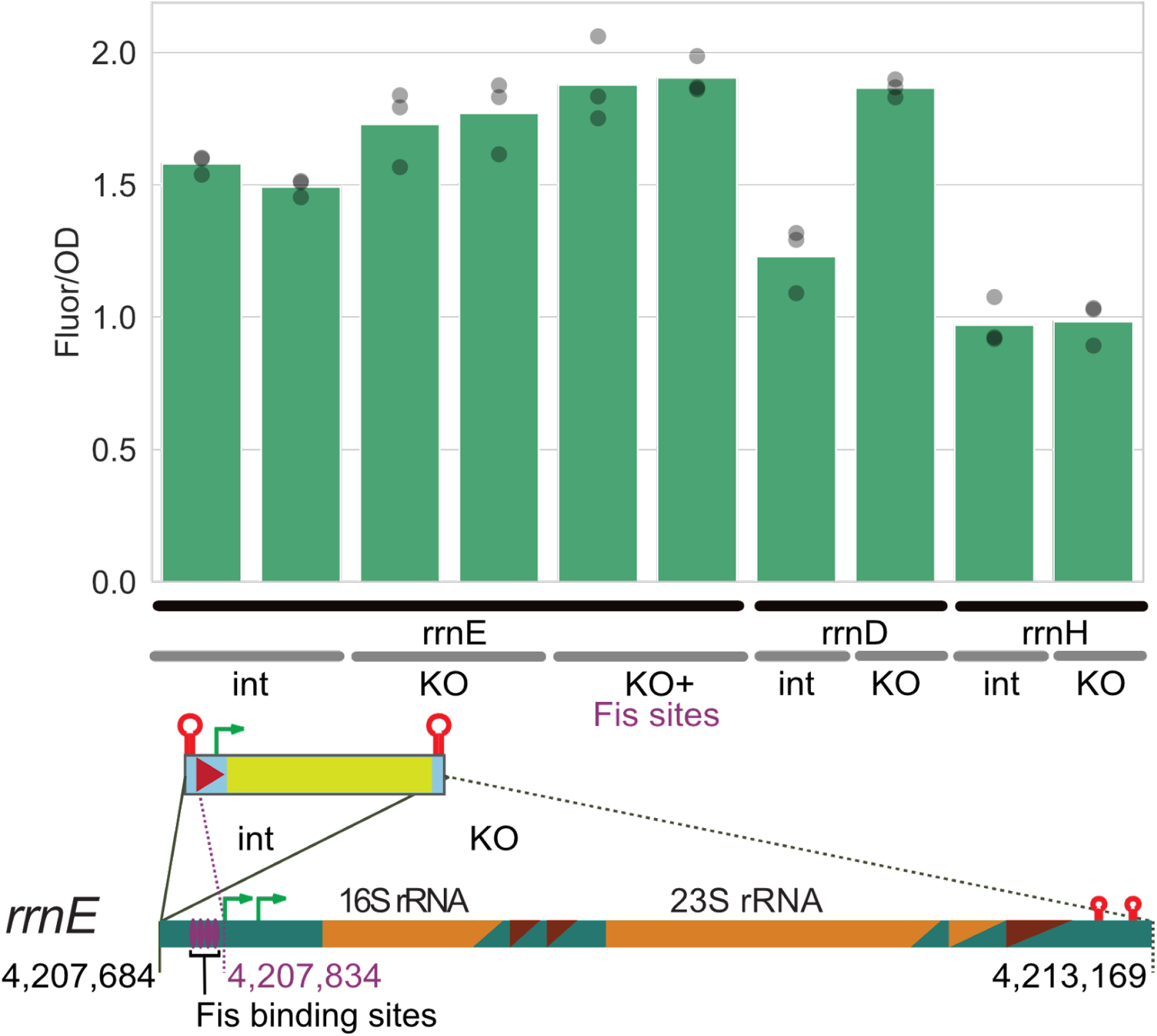
Ribosomal RNA operons are dispensable for regional high transcriptional propensity. The reporter was either integrated upstream (int) of the promoters of ribosomal RNA operons (*rrnE, rrnD, rrnH*) or replacing the entire operon and regulatory elements from the same site upstream of the promoter until after the ribosomal operon terminators (KO), as indicated in the diagram. A KO at *rrnE* that retains the Fis binding sites upstream of the reporter was also tested (KO+; dashed purple line indicates the upstream integration site). Reporter fluorescence per OD 450 was measured in three independent microplate reader experiments. Integration coordinates are for the U00096.3 complete MG1655 genome, the current EcoCyc standard. Cases where two bars are shown for a given genotype indicate two independently constructed clonal lineages; for each bar, the geometric mean of three biological replicates is shown, with the individual replicates as gray points.

Although the *rrn* KOs generally had little to no effect on reporter expression, we noticed that the *rrnD* KO in particular had higher reporter expression compared to the *rrnD* int (*rrnD* upstream integration). To test whether the *rrnD* KO caused a local or general increase in mNG expression, we replaced *rrnD* with a kanamycin resistance marker in strains with mNG integrated into different locations. The *rrnD* KO caused similar increases in reporter fluorescence in all of the sites we tested, whereas two other *rrn* deletions (*rrnA* and *rrnC*) did not (**Supplementary Fig. 8**). However, the *rrnD* KO did not result in an increase of mCherry fluorescence (**Supplementary Fig. 9**). It is possible that removal of *rrnD* has a particularly strong effect on other high transcriptional propensity sites (all three sites considered in **Supplementary Figure 8** are high propensity and fairly close to other *rrn* operons), although the reason for the outsized effect of *rrnD* deletion relative to deletions of other *rrn* operons remains unclear.

### Transcriptionally insulated reporters are affected by strongly expressed genes in a orientation specific manner

Consistent with previous studies, we observed that expression from the mNG reporter integrated directly adjacent to the highly-expressed *rrn*E operon varied depending on the relative gene orientation (3, 26). By testing all possible orientations of the well-insulated mNG reporter integrated at *rrn*E-adjacent sites, we confirmed that sites downstream of this highly-expressed operon show lower transcription compared to upstream integrations (**Fig. 5**). Reporter integrations transcribed in the same direction (tandem orientation) with respect to the *rrnE* operon, whether upstream or downstream, were expressed more highly than those in the divergent or convergent orientation. Our findings are generally consistent with a previous study demonstrating high transcriptional interference between convergently oriented fluorescent protein genes encoded on a plasmid and assessed in a microplate reader (26), and are fully consistent with expectations based on a buildup of negative supercoiling upstream of a highly expressed operon, and positive supercoiling downstream of it. Unlike virtually all other cases considered through our work, in the case of the directional integrations shown in **Figure 5**, we observed a discrepancy in rank ordering between the plate reader-based bulk fluorescence data and flow cytometry data (**Supplementary Fig. 10**); in the latter case, the lowest expression was observed for the Downstream-Tandem rather than Downstream-Convergent orientation. While the vast majority of measurements were concordant between the methods, for an unidentified reason, there was a discrepancy between fluorescent measurements in the microplate reader and flow cytometer in the isolated case of strains with mNG integrated downstream of *rrnE* in the convergent orientation (**Supplementary Fig. 11** and **Supplementary Table 1**). Therefore, we repeated microplate measurements for a subset of specific strains showing a discrepancy between the approaches (**Supplementary Fig. 11**). This repeated trial confirmed a strong reduction in fluorescence of the mNG reporter integrated downstream of *rrnE* in the convergent orientation when measured by microplate reader (**Supplementary Fig. 4**), supporting the conclusions drawn from **Figure 5**. The remaining divergence between the measurement modalities is possibly due to some specific sensitivity of the *rrn* regions to the harvesting conditions required for flow cytometry measurements.

**Figure 5:**
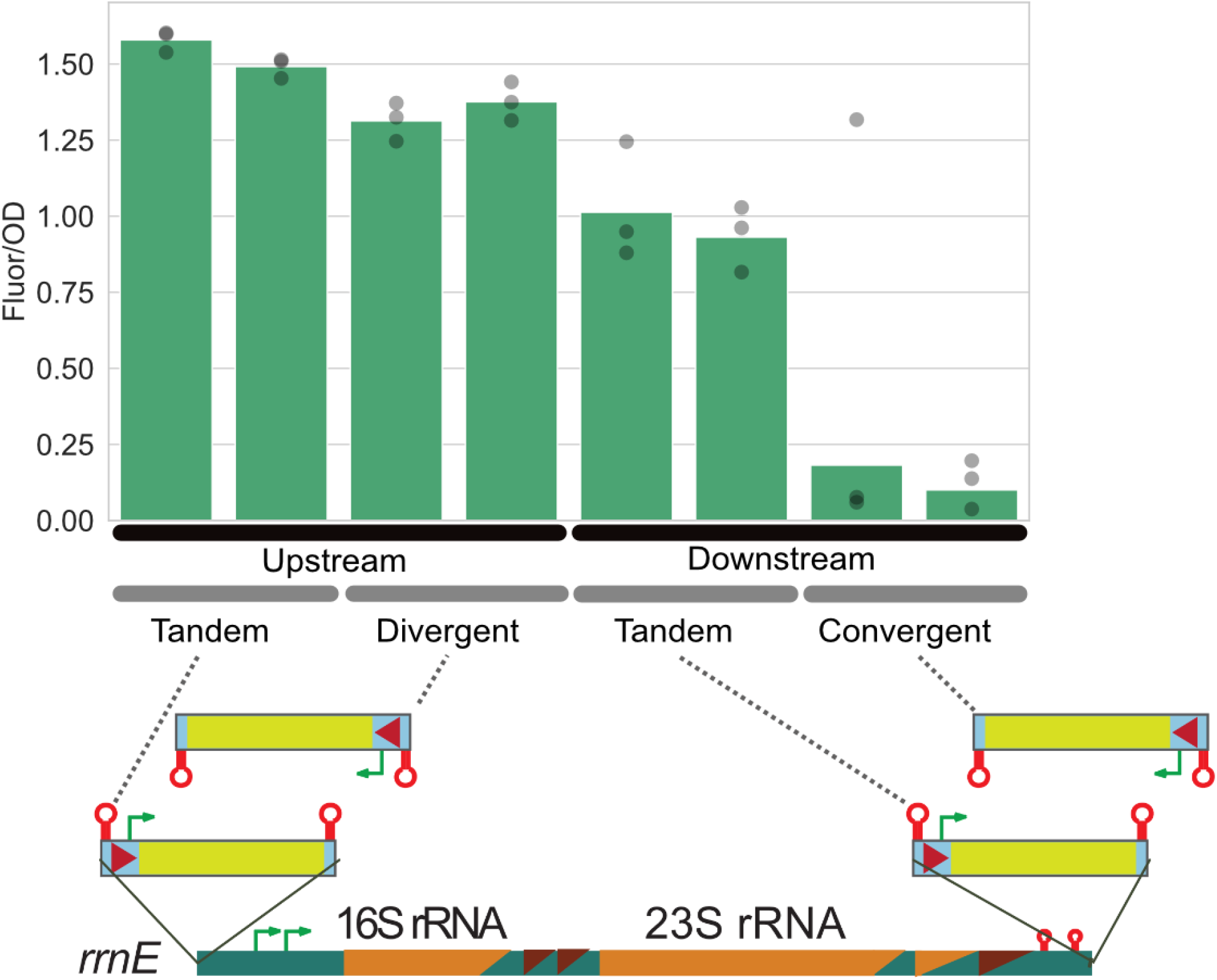
Ribosomal RNA operons affect local direction-dependent gene expression. The reporter construct was integrated into the genome either upstream of the *rrnE* promoters or downstream of the rrnE terminators in all possible orientations as indicated in the diagram. Reporter fluorescence per OD 450 nm measured in three independent microplate reader experiments. The green bar corresponds to the geometric mean of the three replicates (grey circles). Side-by-side independent lineages were derived from separate colonies in the initial reporter integration (shown as separate bars). We observed a single outlier in one of the technical replicates from a strain with the reporter integrated into the convergent orientation with respect to *rrnE*, but it does not substantively affect our conclusions (Although this is the apparent outlier in microplate replicate 1, shown in **Supplementary Fig. 7**, it is still included in the averaging and data analysis shown; see also **Supplementary Fig. 5**).

## Discussion

We investigated the contribution of the chromosomal features with the strongest correlations to position-dependent transcriptional propensity. As local GC content, proximity to rrn, and binding of the NAPs H-NS and Fis co-vary with transcriptional propensity, they were our top candidates for chromosomal features that may be functionally responsible for position-dependent expression variation (1). In general, we designed a strategy to fix a reporter position and sequence while varying the genetic context around it in order to test the contribution of chromosomal features on gene expression. The transcriptional propensity from our previous study was broadly consistent with both the RNA level and fluorescent protein expression level measured from targeted integrations (**Fig. S1-S2**), which allowed us to report mNG fluorescence at the population and single-cell level to assess reporter expression due to position-dependent transcription effects. We note that in contrast to the majority of recent work on the effects of sequence contexts on transcription, which focus on the regulatory sequences immediately upstream of a gene (the promoter region and region 100-200 bp upstream of the transcription start site), here we have kept a fixed sequence in the window 162 bp upstream of the transcription start site (and 852 bp downstream of the end of our reporter ORF), and instead focused on the longer-range impacts of genetic context. Our data demonstrate that, in parallel to the extensively studied fundamental effects of regulatory sequences (6, 34–38), genetic context further than 162 bp from the transcription start site has a profound effect on transcription level, and must be considered both in efforts to model transcription of natural genes, and in the design of synthetic biology constructs.

### Transcriptional propensity affects natural promoters of varying strengths

In our previous work on empirical transcriptional propensity mapping, we used the TetO1 promoter to drive mNG expression and generate the map. As are many promoters used in synthetic biology, the TetO1 promoter is relatively strong, ranking in the 86th percentile of *E. coli* transcriptional output. We wondered whether weaker promoters would be similarly affected by genetic context. Competitive interplay between H-NS and RNAP may lead to greater position-dependent H-NS silencing effects on genes that are naturally more weakly expressed (39). On the other hand, context-dependent silencing of weak promoters may be less apparent, as silencing a weak promoter would result in a relatively small decrease in signal compared to silencing a strong promoter assuming some basal level of transcriptional activity (40). By swapping only the reporter promoter to one of three additional sequences with a range of strengths at three chromosomal sites (*eaeH, yagF*, and *yafT*) that are close to each other (and therefore have comparable DNA copy number) but have different transcriptional propensity (**Fig. 1**), we demonstrate that natural promoters are subject to similar context-dependent expression variation (**Fig. 1**). The same conclusion has also been drawn from many thousands of promoters tested at three different genomic loci (41). Only our weakest promoter was unaffected by the genomic context, which is consistent with a context-independent transcriptional activity baseline. Indeed, the detection of RNA that is antisense to genes or intergenic in high-coverage RNA-seq experiments, which may be deleterious to cells, is presumably a result of “leaky” transcription in at least some cells (40, 42, 43) and may occur at some level independent of any transcriptional propensity effects or regional silencing.

### Extreme GC content flanking regions overrides regional transcriptional propensity

H-NS binds to and silences sites of low GC content (17–20). In our high-resolution transcriptional mapping study, we found that the mNG reporter (54% GC content) became silenced when it integrated into H-NS bound regions at different sites across the genome (1). Here, we report that synthetic context flanks composed of 35% GC can strongly silence transcription even in an otherwise high transcriptional propensity site, and that 65% GC flanks can partially relieve the repression of a low transcriptional propensity site (**Fig. 2**). How can flanking DNA have such a dramatic effect on gene expression? H-NS may be forming silencing filaments from high-affinity low GC sites flanking the reporter (44) to invade and silence the reporter itself. Previous work has also shown that H-NS and associated proteins StpA and Hha can form bridge filaments leading to early termination of RNAP elongation (21, 45). These mechanisms may explain the reduced transcriptional propensity for an otherwise highly-expressed and GC neutral (54% GC) reporter when integrated into low GC content genomic regions bound by H-NS, such as near wbbH.

We note that in our sampling of different promoters and placement contexts, the mNG concentration in individual cells within a population always shows a unimodal shift in cell-level expression (**Fig. 1** and **Supplementary Fig. 3**). This indicates that the drivers of transcriptional propensity that are being modulated here (likely H-NS silencing) are not stochastic events that are biased towards a higher frequency due to context, but rather affect all cells in a tunable manner. Furthermore, we never observed the appearance of a secondary population of cells with a distinct fluorescence level in any of the strains tested in this study. Whether the reporter fluorescence was decreased by low flanking GC content or by transcriptional interference from *rrnE*, the cells fluorescence level shifted as a unimodal population. We emphasize that the accumulation of cells with low mNG concentration represented in the flow cytometry violin plots in this article is simply the result of assigning cells without detectable fluorescence above the median autofluoresce level (from reporter-free cells) to a minimum fluorescence level across experiments. This assignment clearly indicates the point at which cells do not have detectable fluorescence, while maintaining the median and quantile values of the population level fluorescence.

A previous study has shown that altering the GC content of a gene in a synthetic operon can affect the expression of an adjacent gene in the same operon (46), consistent with the non-coding GC content effects presented here. These observations may be a result of H-NS binding in addition to the changes in RNA folding energy reported in that study. Non-coding, low GC content can cause silencing of a neutral-GC reporter in the chromosome and on a plasmid. This silencing can be evolutionarily favored, at least on a plasmid, by imposing a lower RNA/protein production burden on the cell (47). Taken together, these findings suggest that low GC content DNA can be horizontally transferred and maintained at a relatively low cost and a low level of expression of potentially toxic gene products (even if short stretches of neutral GC content are included), which represents a reservoir of latent genetic material shared between compatible bacteria, as has been extensively discussed in the context of xenogeneic silencers (48).

### Transcriptional propensity affects maximal induction but not induction kinetics

Although we observed a dramatic reduction in mNG level from reporters integrated into low GC content synthetic contexts, there were no major differences in the time required for maximal induction of the reporter (**Fig. 3**). The silencing effect of low GC content flanks was also completely resistant to extended induction times. In other words, transcriptional activity did not cause further de-repression of mNG expression in low GC contexts, unless over short periods of tens of minutes or less. Thus, it appears that the context effects that we have termed transcriptional propensity act mainly on the equilibrium rate of transcription achieved. The induction data also show that our previous *E. coli* random integration library cells, used to produce our high-resolution transcriptional propensity map, should have achieved the maximal expression level achievable given the genetic context of each reporter.

### The high transcriptional propensity near ribosomal RNA operons is a genomic context feature that does not require the operon or its promoter

Ribosomal RNA operons (*rrn*) are another key structural element of the *E. coli* chromosome as they form some chromosome interacting domain boundaries (49). *rrn* can also form nucleolus-like zones in the nucleoid where they colocalize with foci of actively transcribing RNAP (24, 50, 51), with focus formation dependent upon the nucleoid-associated protein HU (52). Directly around *rrn* there are broad transcriptional propensity peaks, suggesting that these chromosomal regions may be brought into or are affected by transcription factories (1). In order to test the functional contribution of the *rrn* themselves to the high transcriptional propensity around them, we made targeted reporter integrations either upstream of, or replacing, the *rrn*. Replacement of the *rrn* did not cause a decrease in reporter fluorescence compared to the corresponding upstream integration (**Fig. 4**). The *rrn* KO removes the entire transcription unit and regulatory features, including important features required for *rrn* colocalization, such as the promoters (24). Thus, the *rrn* operons and their *cis* regulatory regions do not set the transcriptional propensity in the genomic region where these operons are found, but rather, the neighboring context must have evolved to enhance expression from the regions of *rrn*. Future work may determine the effect of physical localization of chromosomal regions and of potentially unidentified sequence features on local transcriptional propensity. Unexpectedly, some of the *rrn* replacements with the reporter construct in fact caused an increase in reporter fluorescence. The effect was especially pronounced for the *rrnD* replacement. We found that *rrnD* KO by a Kanamycin resistance cassette also caused an increase in reporter fluorescence that was integrated into other sites around the genome (**Supplementary Fig. 8**). We could not attribute this change in reporter fluorescence to a change in cell size or growth rate (**Supplementary Fig. 7**), nor was mCherry fluorescence from a gene present in our base strain affected by the *rrnD* KO (**Supplementary Fig. 9**). A change in relative expression level for a cluster of translation factors proximal to *rrnD* is a possible explanation for these observations. Testing of additional reporter integration sites in the WT and *rrnD* KO context might determine whether some genomic regions are in competition with *rrnD* for limiting expression resources, such as RNAP.

### Supercoiling interference extends past transcript boundaries and transcriptional terminators

Finally, we also confirmed interference effects that have previously been observed on gene expression when encoded downstream of an induced gene, regardless of the downstream gene orientation (3). In order to test the interference of a strongly expressing gene on the mNG reporter, which is insulated by very strong well-characterized terminators (53), we integrated the mNG reporter in all possible orientations around the highly expressed *rrnE* operon, without disrupting the promoter, associated Fis binding sites, or the operon terminators. Consistent with the previous studies, we find that the strongest interference effects on reporter expression when the reporter is integrated downstream of the strong transcriptional unit *rrnE*, especially in the convergent orientation (**Fig. 5**). In general, our results regarding the effects of supercoiling interference for a reporter integrated into the genome are in agreement with previous results in which reporter genes were encoded on a plasmid (26). There may be a broader range of mNG fluorescence in strains with the reporter integrated in the convergent orientation (**Supplementary Fig. 10, Supplementary Fig. 4**), which might be consistent with an observation of increased expression “noise” for a reporter integrated into *rrn* operons in a previous study (16). However, expression noise from a reporter integrated at a site within an *rrn* may also be due to reporter loss by recombination and a competitive advantage for cells with 7 functional *rrn*, especially in rapid growth conditions.

Repressive effects have also been observed for genes in the divergent orientation even over relatively long (>2 kb) intervening lengths, but the length limit of this effect is not known (30). *In vitro* evidence suggests that DNA gyrase can relieve the effects of divergent supercoiling competition (26). In general, our findings are consistent with divergent supercoiling competition, but the effect size is small (**Fig. 5**). Genetic interference effects can be further complicated by transcription units without completely insulating terminators (54). Subtleties in the exact genetic construct design and spacing are likely responsible for variations in effect size reported between and within studies. Future work on a greater diversity of genetic construct designs using a similar induction matrix as Yeung et al. (26) and single-cell analysis may be an important, albeit intensive, undertaking to fully understand competition between neighboring transcriptional units. Expanded mechanistic insights into context effects will allow utilization of genetic context as a modulating control mechanism in synthetic biology (55). Regardless of the application, genetic context should be taken into consideration. This has been demonstrated by a recent effort to standardize genomic integration designs, where sequence-controlled expression level and integration frequency were variable depending on the landing pad position (56). The findings presented here demonstrate not only the importance of such endeavors, but the long range of context effects that must be considered, and the extent to which synthetic flanking sequences can overcome natural context effects on the bacterial chromosome.

## Conclusion

Local and regional genetic contexts have a strong influence on expression from a transcriptionally insulated reporter expressed from both strong and weak promoters. These context effects are key considerations that should be taken in parallel with other genetic construct design choices like promoter, RBS and terminator for plasmid cloning and for integration into genomes. Genetic context manipulation is also a unique tool in the genetic toolbox for tuning gene expression without altering the coding or regulatory element sequence for genes in synthetic metabolic pathways or genetic circuits.

## Acknowledgements

We are grateful to Rebecca Scholz for critical feedback on the manuscript. This work was supported by NIH R03-AI130610 (to P.L.F.), and R35-GM128637 (to P.L.F.). S.A.S. was additionally supported by the NIH Cellular and Molecular Biology Training Grant (T32-GM007315). C.D.L. was additionally supported by the NIH Predoctoral Training Program in Genetics (T32-GM007544).

