## Supplementary Information for "Genetic context effects can override canonical *cis* regulatory elements in *Escherichia coli*"

### Supplementary Information for “Genetic context effects can over-ride canonical cis regulatory elements in Escherichia coli”

Scott A. Scholz[1,†], Chase D. Lindeboom[1], and Peter L. Freddolino[1,2,*]

1 Department of Biological Chemistry, University of Michigan Medical School, Ann Arbor, MI, USA

2 Department of Computational Medicine and Bioinformatics, University of Michigan Medical School, Ann Arbor, MI, USA

† Current Address: Max Planck Institute for Terrestrial Microbiology, Marburg, Germany

#### Supplementary Text

##### Supplementary Text 1: Calibration of RNA and protein readouts of transcriptional propensity

In order to compare to our previously generated high-resolution transcriptional propensity mapping data, we used the same reporter construct for the experiments presented here, with targeted integrations into sites representing a range of transcriptional propensities. As the majority of the new integration locations considered here did not match an integration from the original library measurements of (1), we predicted the transcriptional propensity of each newly integrated site based on the original library measurement by extracting all original data points within a 5 kilobase window of the new insert location, and fitting a LOESS model to the data points in that region (using 0.2 for the span parameter); the LOESS-predicted propensity of the new point was then used.

We observed that strain fluorescence produced by this reporter upon targeted integration at new sites is generally consistent with our high-resolution transcriptional propensity mapping data (1) (**Supplementary Fig. 1**), whether measured at the population level or at the single cell level. In addition, we observed that for several tested sites, the transcriptional propensity, RNA levels, and protein levels measured at both bulk and single-cell levels were highly correlated with slopes near 1 (**Supplementary Fig. 2**); thus protein level readouts for this reporter are a faithful proxy for transcript levels. This was expected for an unchanging RBS and reporter sequence and because a substantial proportion of protein level variation in *E. coli* is regulated at the transcriptional level (57, 58). Although these data do not rule out the possibility of genome-position dependent translation effects, they show that we were able to track fluorescence as a readout for genetic context dependent transcription variation for the specific loci examined herein. We note also that while in our prior work we considered both raw and dosage-adjusted transcriptional propensities, the latter of which accounts for DNA copy number variations over the length of the chromosome arising due to DNA replication, in the present work we focus solely on the original (unadjusted) transcriptional propensity due to the different growth conditions employed here (most notably, frequent use of wells in a 96 well plate grown in a plate reader), which may alter the dosages present over the length of the genome.

We remark here that two sites shown in **Supplementary Figure 1**, *yqjI* and *yagF*, appear to be outliers from the overall agreement observed between transcriptional propensity and fluorescence measurements. Our targeted integrations at *yqjI* and *yagF* showed substantially higher and lower expression levels, respectively, than would be expected based on the previously measured transcriptional propensities at those regions. There are several possible explanations for this observation. As the original transcriptional propensity signal represents an average over several (at least three) sites on a 500 bp window, any single integration may not necessarily display the expected expression level if short length scale variations exist in the propensity; we have not studied the potential for such high-frequency effects in detail, and neither the *yqjI* nor the *yagF* sites used here were in the original library. Another possibility is that the neglect of gene dosage compensation in the present analysis is partly responsible for the discrepancy. It is also notable that of the sites shown in **Supplementary Figure 1**, *yqjI* and *yagF* are the only two for which our integrations were inside of open reading frames, and the resulting unexpected expression levels may result due to transcriptional interference from the native genes. It is also possible that these sites represent artifactual outliers in the original transcriptional propensity map, or that minor differences in physiological conditions between the original and present experiments are sufficient to alter the propensities at those locations. Although we did not observe a global bias in DNA barcode counts with local genomic GC content, it is possible that at the two most extreme GC-rich regions such as those surrounding the *yagF* (also tested in this work) and *nrfG*, a reduced amplification of integrated DNA barcodes, as previously noted (59), lead to outliers in the transcriptional propensity map at those sites (1). While the reasons for the *yqjI* and *yagF* sites acting as outliers are unclear, they are excluded from the fits shown in **Supplementary Fig. 1-2,** which are aimed primarily at calibrating the relationship between various forms of nucleic acid and protein-level measurements of our reporter construct at different sites. Future investigation will allow us to further pinpoint the reason for the unexpected behavior of these particular integration sites. It is also notable that even for the outlier *yagF* site, excellent correlation was observed between the qPCR, flow cytometry, and bulk fluorescence measurements used throughout this work (**Supplementary Fig. 2**, bottom panels), and the observed expression is higher than that of nearby sites with lower transcriptional propensity (**Fig. 1**).

#### Supplementary Figures

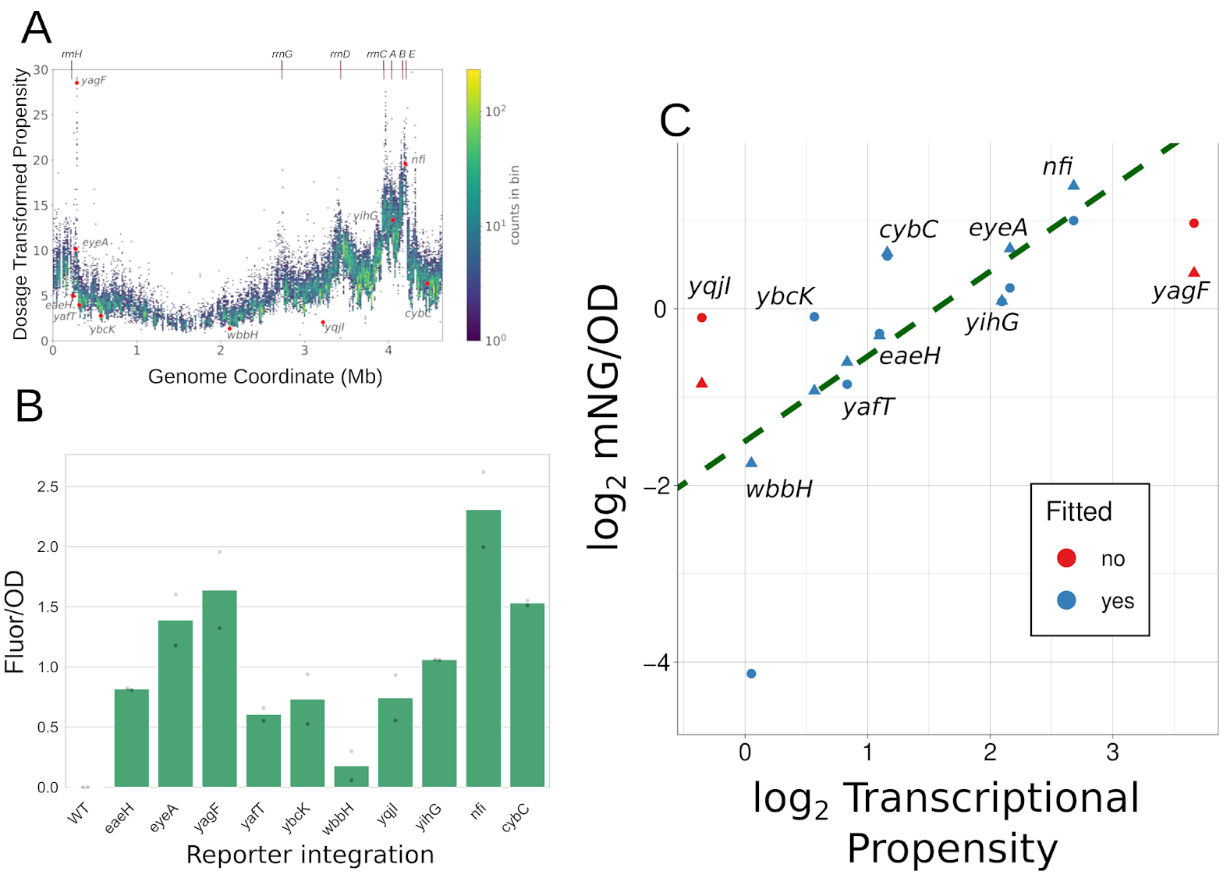


**Supplementary Figure 1: Fluorescence from targeted integrations of a standardized reporter spans a broad range.** A) Locations of all targeted integrations used in this study mapped to a DNA dosage-corrected high-resolution transcriptional propensity map. Transcriptional propensities for the new integrations are predicted based on interpolation from the original library data; see Methods for details. B) Fluorescence per OD 450 from the mNG reporter integrated into sites (site and transcriptional propensity shown in A) chosen to span a broad range of transcriptional propensity. C) Correlation of fluorescence/OD and dosage propensity after removing outlier *yagF* and *yqjI* sites (shown is a robust linear regression; slope: 0.96; standard error on fitted slope: 0.16; p=0.00004); shapes indicate biological replicates from different days. If the outliers are included in the fit the regression still shows a significant but weaker correlation between transcriptional propensity and expression levels (slope: 0.51; standard error: 0.14; p=0.002); the motivation for the exclusion of *yagF* and *yqjI* is discussed in **Supplementary Text 1**.

.

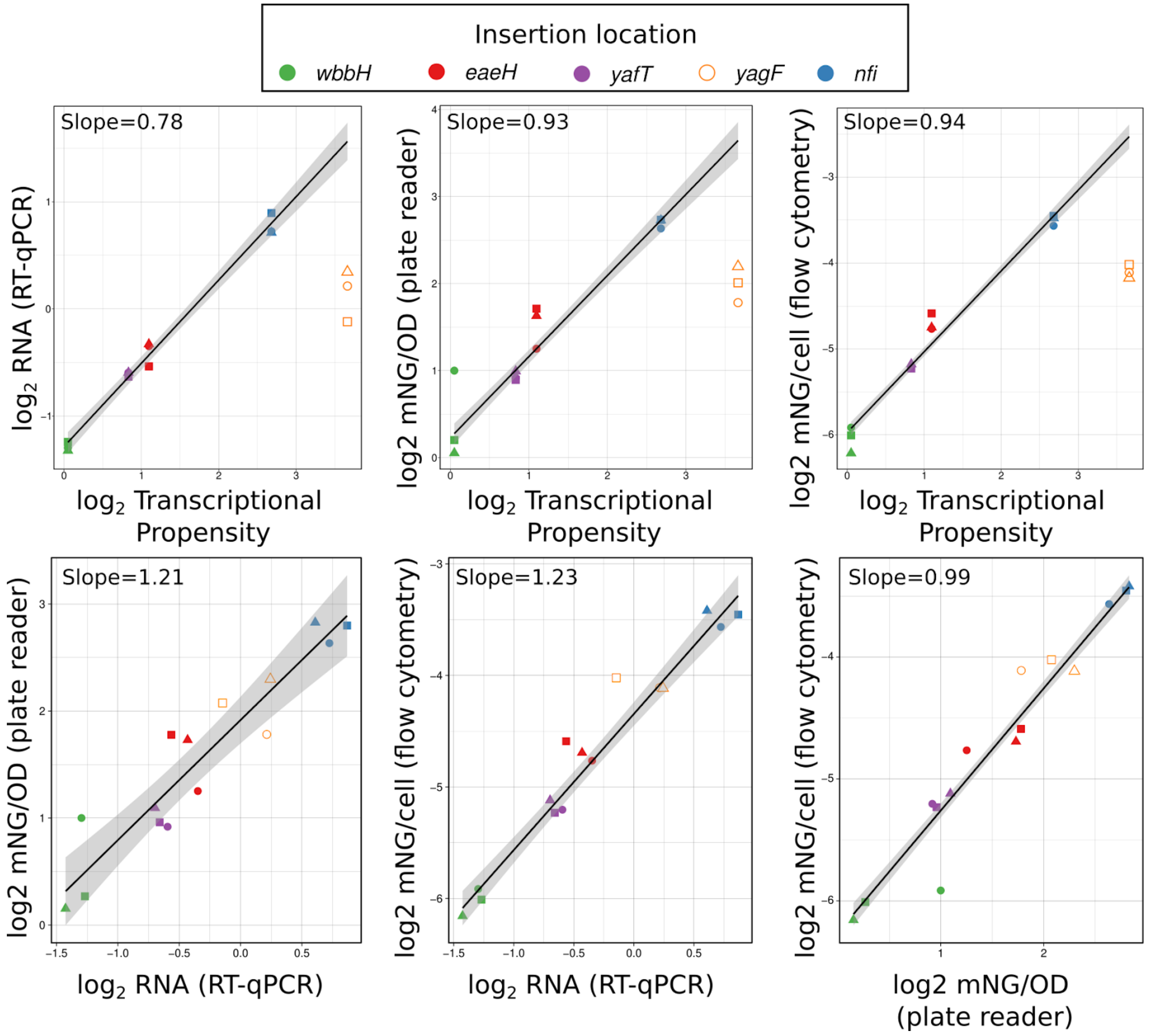


**Supplementary Figure 2**: **Fluorescence is a faithful reporter of transcriptional propensity.** We assessed the correlations between transcriptional propensity, RNA levels (via RT-qPCR), and fluorescent protein production, making use of five different integration sites (all are also shown in **Supplementary Fig. 1**). Colors indicate the integration locations, and differently shaped points are biological replicates from different days. Shaded regions show bootstrap-based 95% confidence intervals for the slope. Top row: comparisons of the correspondence between transcriptional propensity and each of three possible follow up measurements; robust linear regressions are shown after correction for replicate-level effects (obtained using the `rlm` function in R, with a model containing both genotype and biological replicate terms). The *yagF* site is excluded from the fits (and thus shown with points unfilled, as discussed in **Supplementary Text 1**). Bottom row: Correlations between the low-throughput transcriptional propensity proxies considered here.

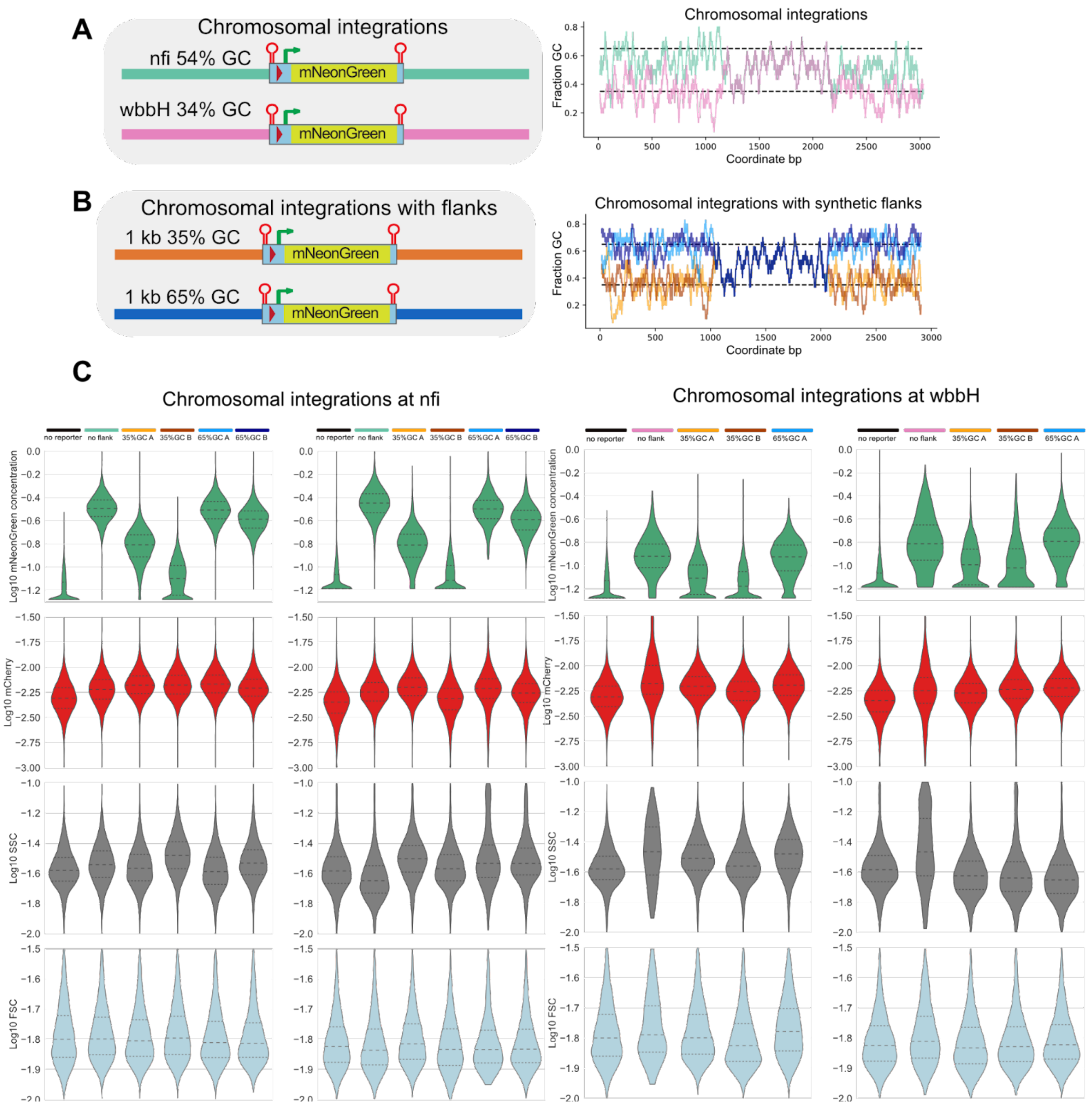


**Supplementary Figure 3**: **Synthetic flanks of varying GC content cause unimodal shifts in mNG concentration distribution of individual cells.** A) Diagram of the mNG reporter integrated into nfi or wbbH locus with a corresponding sliding window 30 bp of average GC content. B) The same diagram as in A for mNG reporters integrated with synthetic flanks. C) Violin plot distributions of flow cytometry measurements of mNeonGreen concentration with two independent replicates shown side-by side. Growth problems precluded measurement of the *wbbH* integration of the 65% GC B reporter alongside the other cases shown above. We also note that the effect of high GC flanks on *wbbH*-adjacent expression appear weaker based on flow cytometry than they do based on plate reader data; the plate reader findings were further confirmed through additional replicates shown in **Supplementary Figure 4**, and thus the most plausible explanation is that the different growth conditions required for flow cytometry have some blunting effect on the activation provided by the 65% GC A reporter at *wbbH*.

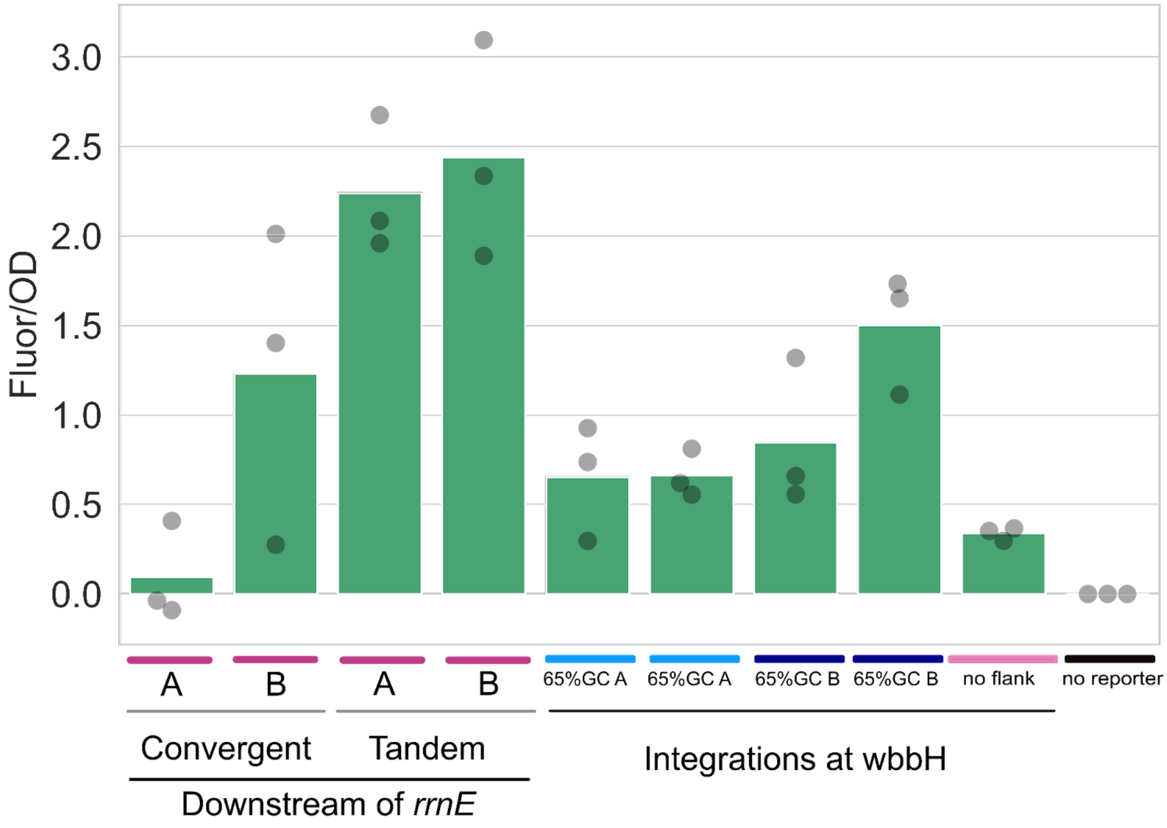


**Supplementary Figure 4: An additional microplate reader assessment of a subset of strains conducted at a different time confirms two different types of genetic context effects, as expected.**

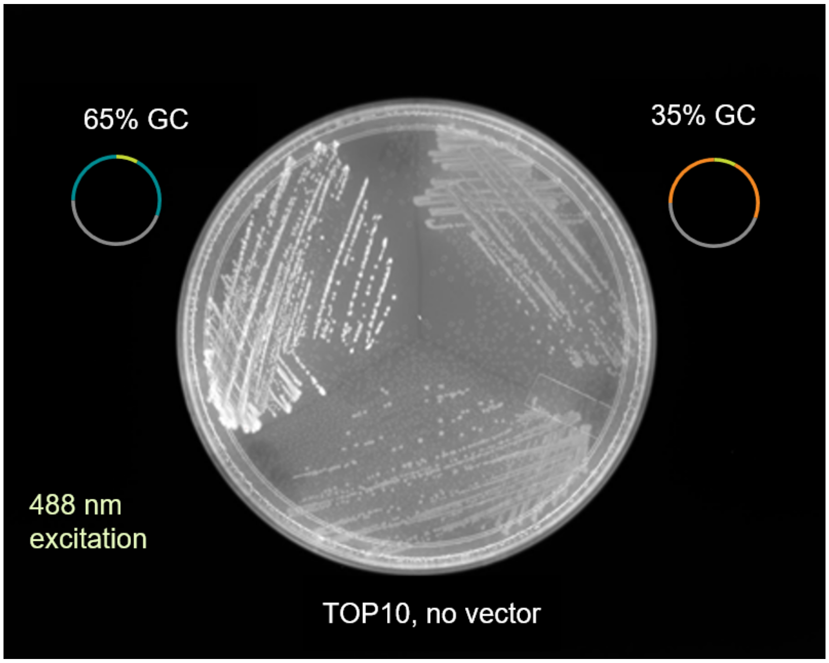


**Supplementary Figure 5: Genetic context affects fluorescence reporter expression on plasmids.** Top10 E. coli cells containing either the plasmid pSAS70 (3.5 kb of synthetic 35% GC content flanking the reporter on either side, represented in orange), pSAS72 (3.5 kb of synthetic 65% GC content flanking the reporter on either side, represented in blue) or without any plasmid were streaked out on an LB plate, grown overnight at 30º C and illuminated in 488 nm wavelength light. Only the reporter surrounded by 65% GC content shows strong fluorescence.

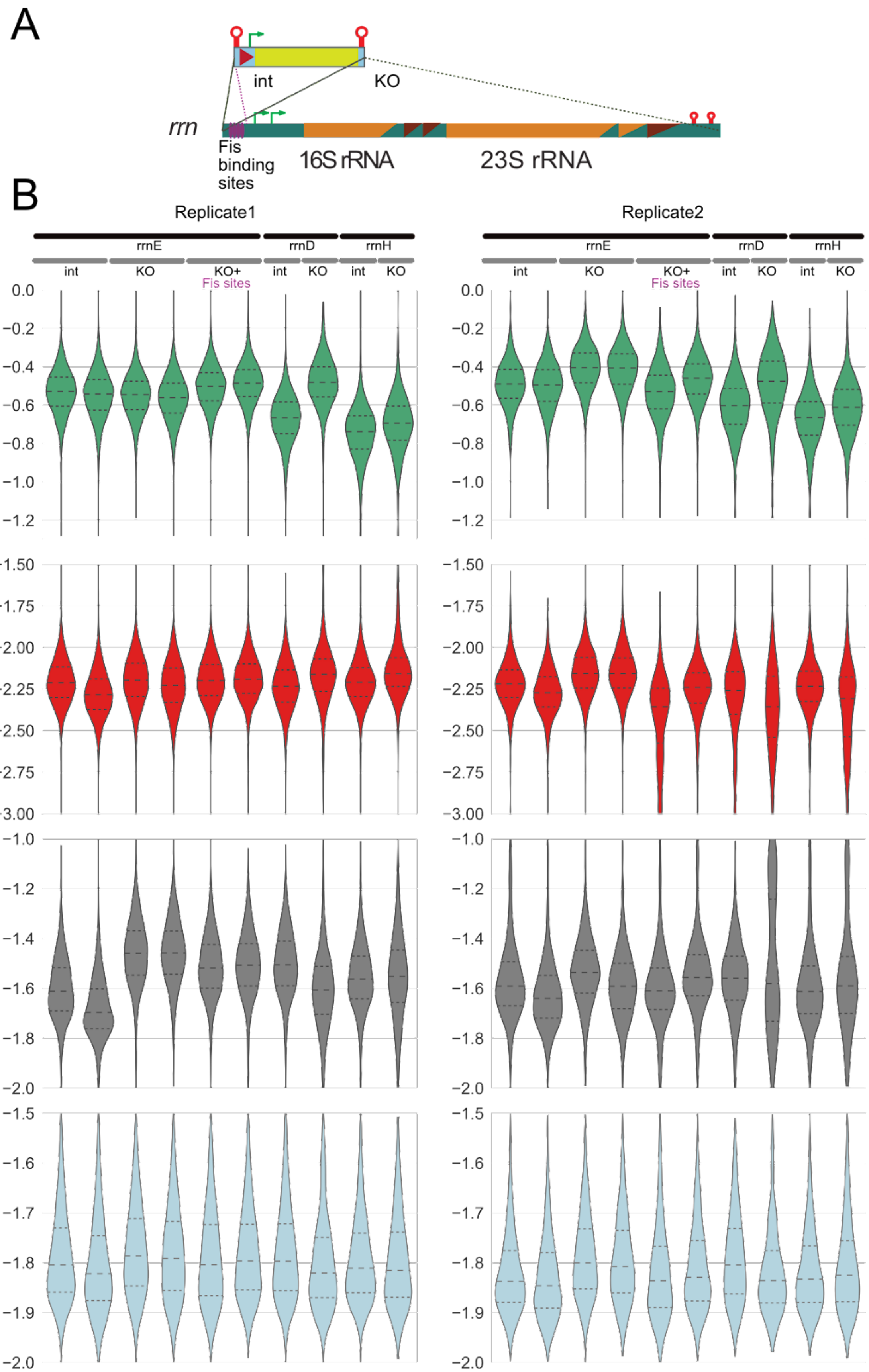
A
int
KO
rrn
Fis binding sites
16S rRNA
23S rRNA
B
Replicate1
Replicate2
rrnE
rrnD
rrnH
int
KO
KO+ Fis sites
Log10 mNeonGreen concentration
Log10 mCherry
Log10 SSC
Log10 FSC

**Supplementary Figure 6: Distribution of mNG concentration in cells with the reporter integrated either at or replacing rrn.** A) diagram of mNG integrations at rrn that insert upstream of the rrn, replace the rrn or replace the rrn while retaining upstream Fis binding sites. B) Violin plot distributions of flow cytometry measurements of mNeonGreen concentration with two independent replicates is shown. Below in red, the distribution of mCherry signal above autofluorescence level of the MG1655 base-strain without mCherry integrated into the genome is shown. The distribution of the side-scatter signal is shown in grey. The distribution of the forward-scatter signal is shown in blue.

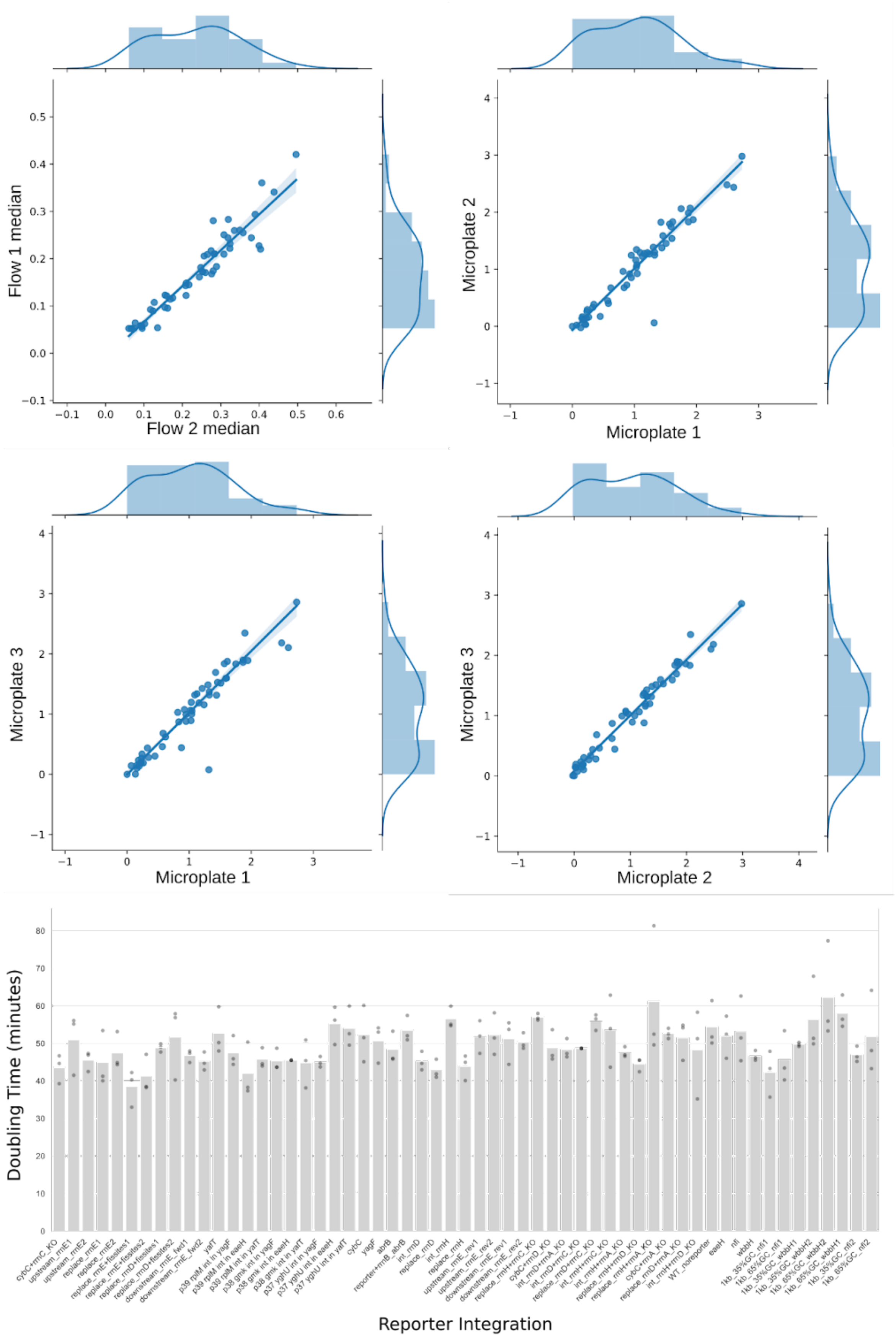

Flow 1 median
Flow 2 median
Microplate 2
Microplate 1
Microplate 3
Microplate 1
Microplate 3
Microplate 2
Doubling Time (minutes)
Reporter Integration

**Supplementary Figure 7: Scatter plots of replicate experiments show that variation in reporter expression is consistent across experiments.** Spearman correlation values are shown in **Supplementary Table 1**. Below, the strain doubling time is shown during log-phase growth for every strain grown simultaneously in the microplate reader for three independent experiments.

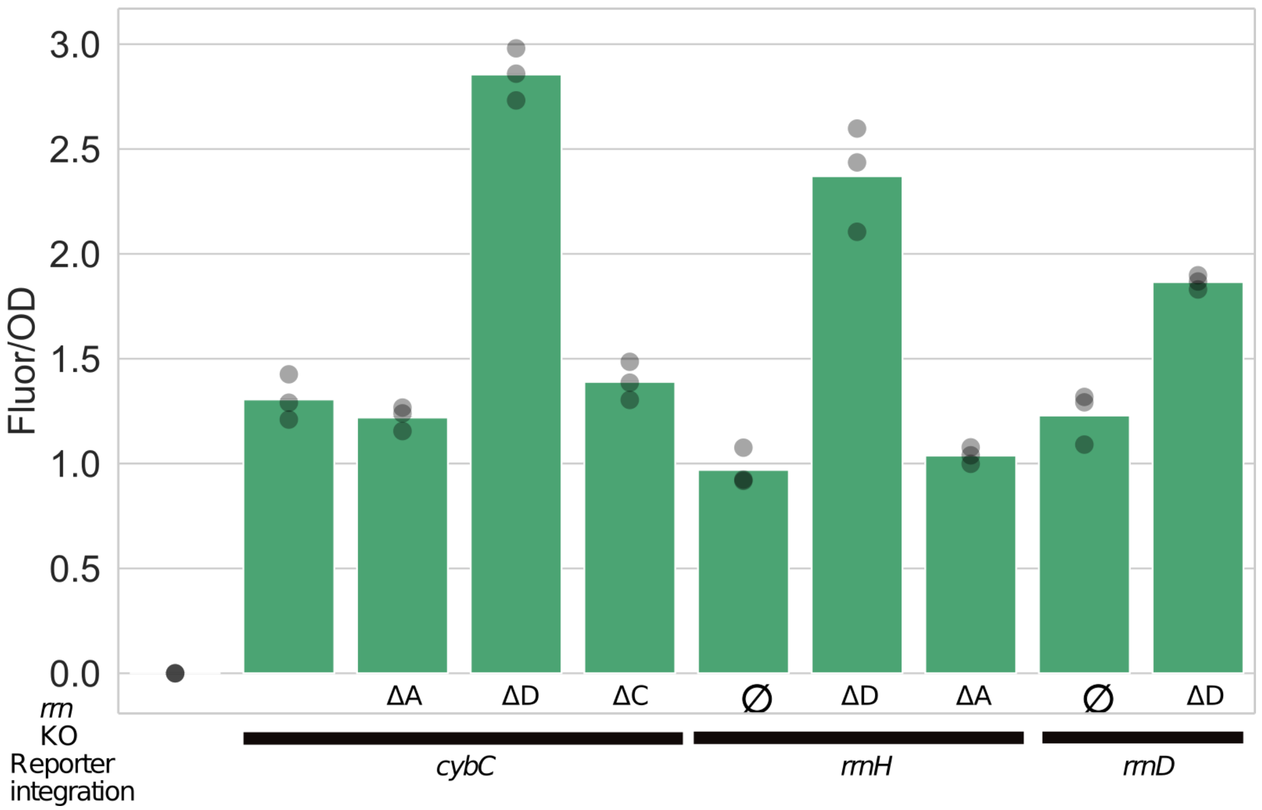


**Supplementary Figure 8: Knockout of *rrnD* causes an increase in fluorescent reporter expression from diverse genetic loci.** The reporter was integrated into different genomic loci with ribosomal RNA operons knocked out (See **Supplementary Figure 1** for a visualization of integration locations). Reporter fluorescence per OD 450nm measured in three independent microplate reader experiments is higher when *rrnD* is knocked out, but not when *rrnA* or *rrnC* are knocked out, regardless of the tested integration sites of the reporter.

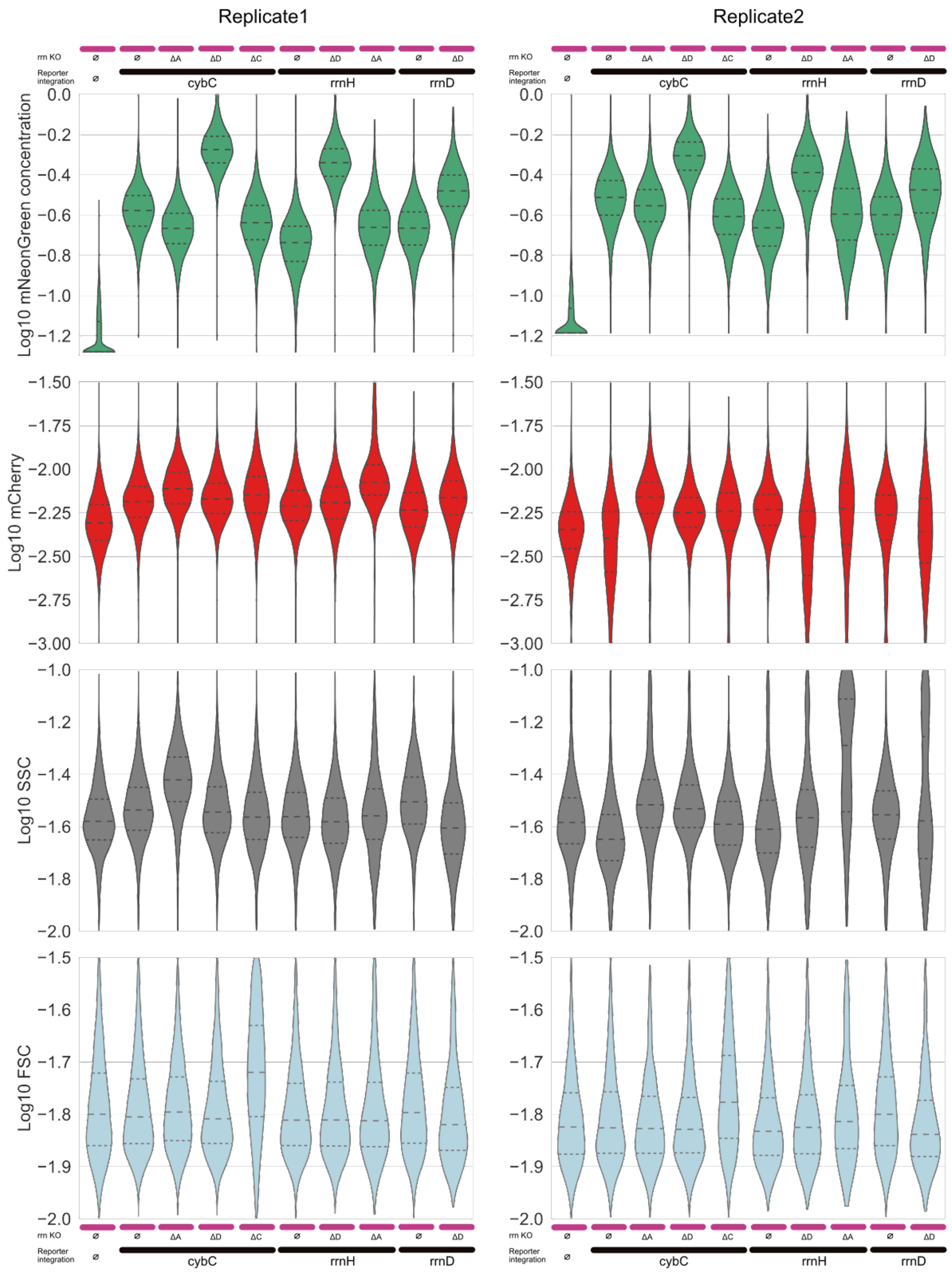


**Supplementary Figure 9**: **Knockout of *rrnD* flow cytometry.** Violin plot distributions of flow cytometry measurements of mNeonGreen concentration with two independent replicates shown side-by side. Below, the distribution of mCherry signal above autofluorescence level of the MG1655 base-strain without mCherry integrated into the genome. Next, the distribution of side-scatter signal and the distribution of forward-scatter signal are plotted.

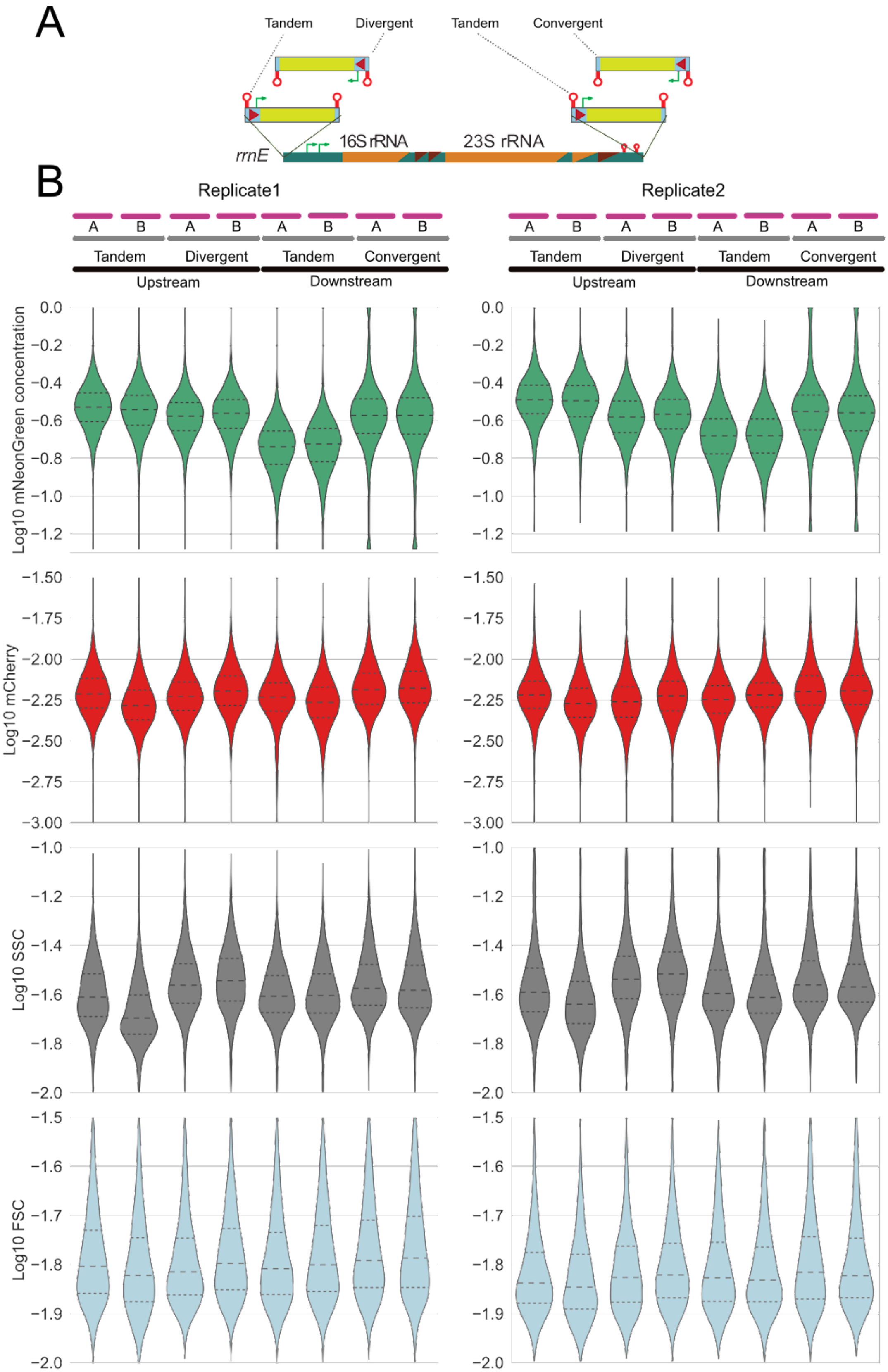


**Supplementary Figure 10: mNG orientation with respect to *rrnE* affects fluorescence of**

**individual cells.** A) Diagram of mNG reporter integration relative to the *rrnE* operon. B) Violin plot distributions of flow cytometry measurements of mNeonGreen concentration with two independent replicates shown. Below, the distribution of mCherry signal above autofluorescence level of the MG1655 base-strain without mCherry integrated into the genome is shown followed by the distribution of side-scatter signal and the distribution of forward-scatter signal.

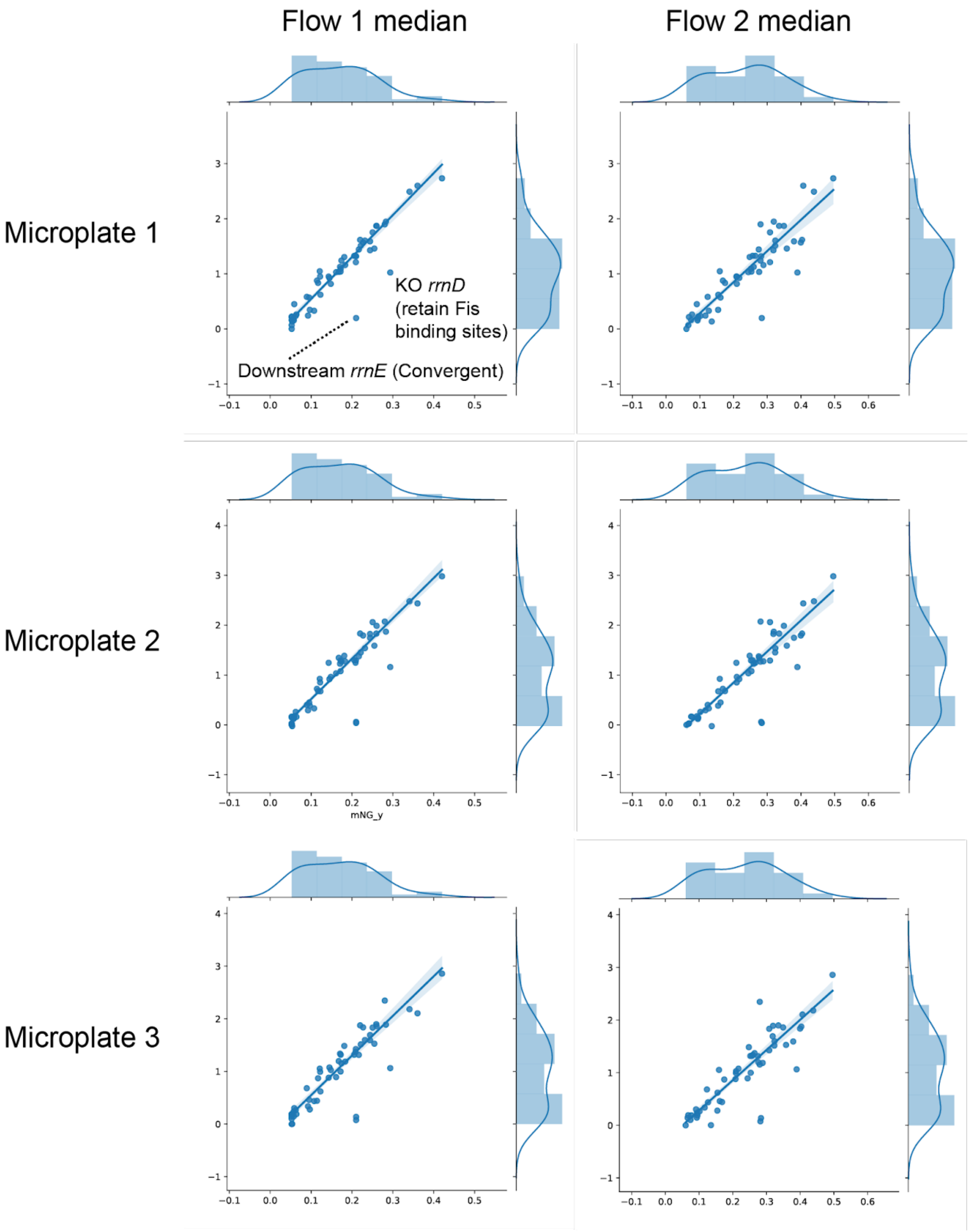


**Supplementary Figure 11: Population level fluorescence levels as measured by microplate reader and flow cytometry are in good agreement.** The median normalized fluorescence of individual cells calculated from flow cytometry data was compared to

Fluorescence/OD 450 for every strain included in all experiments (52 independently constructed strains); definitions match those shown in **FIgure 2** and **Supplementary Figure 3**. In the first panel, labels indicate the two apparent strains in which fluorescence concentration from flow cytometry is not in good agreement with fluorescence/OD 450 from microplate reader assessment. Although the *rrnD* KO (retain Fis sites) strain was not discussed, it was included for completeness, as it was measured in all shown experiments. Spearman correlation values are shown in **Supplementary Table 1**.

#### Supplementary Tables

| | Microplate1 | Microplate2 | Microplate3 | Flow1 | Flow2 |
|---|---|---|---|---|---|
| Microplate1 | 1.00 | 0.95 | 0.94 | 0.92 | 0.89 |
| Microplate2 | | 1.00 | 0.98 | 0.88 | 0.86 |
| Microplate3 | | | 1.00 | 0.87 | 0.85 |
| Flow1 | | | | 1.00 | 0.96 |
| Flow2 | | | | | 1.00 |

**Supplementary Table 1:** The median normalized fluorescence of individual cells calculated from flow cytometry data was compared to Fluorescence/OD 450 nm measured in the microplate reader for every strain included in all experiments (52 independently constructed strains). The Spearman correlation is shown for each comparison, corresponding to **Supplementary Fig. 11** and **7.**

#### Supplementary Data Files

**Supplementary Data 1**: Genbank-formatted sequences of plasmids containing the four high/low GC contexts used in **Figure 2**.

**Supplementary Data 2**: Description of all strains included in this study including reporter integration coordinate.

**Supplementary Data 3**: Primers used for production of PCR fragments used for lambda-red targeted genomic integrations.
